# Multi-Channel Whole-Head OPM-MEG: Helmet Design and a Comparison with a Conventional System

**DOI:** 10.1101/2020.03.12.989129

**Authors:** Ryan M. Hill, Elena Boto, Molly Rea, Niall Holmes, James Leggett, Laurence A. Coles, Manolis Papastavrou, Sarah Everton, Benjamin A.E. Hunt, Dominic Sims, James Osborne, Vishal Shah, Richard Bowtell, Matthew J. Brookes

**Author notes:** **Correspondence to:** Mr. Ryan M. Hill, Sir Peter Mansfield Imaging Centre, School of Physics and Astronomy, University of Nottingham, University Park, Nottingham NG7 2RD, United Kingdom.

## Abstract

Magnetoencephalography (MEG) is a powerful technique for functional neuroimaging, offering a non-invasive window on brain electrophysiology. MEG systems have traditionally been based on cryogenic sensors which detect the small extracranial magnetic fields generated by synchronised current in neuronal assemblies, however such systems have fundamental limitations. In recent years quantum-enabled devices, called optically-pumped magnetometers (OPMs), have promised to lift those restrictions, offering an adaptable, motion-robust MEG device, with improved data quality, at reduced cost. However, OPM-MEG remains a nascent technology, and whilst viable systems exist, most employ small numbers of sensors sited above targeted brain regions. Here, building on previous work, we construct a wearable OPM-MEG system with ‘whole-head’ coverage based upon commercially available OPMs, and test its capabilities to measure alpha, beta and gamma oscillations. We design two methods for OPM mounting; a flexible (EEG-like) cap and rigid (additively-manufactured) helmet. Whilst both designs allow for high quality data to be collected, we argue that the rigid helmet offers a more robust option with significant advantages for reconstruction of field data into 3D images of changes in neuronal current. Using repeat measurements in two participants, we show signal detection for our device to be highly robust. Moreover, via application of source-space modelling, we show that, despite having 5 times fewer sensors, our system exhibits comparable performance to an established cryogenic MEG device. While significant challenges still remain, these developments provide further evidence that OPM-MEG is likely to facilitate a step change for functional neuroimaging.

**HIGHLIGHTS:**

- A 49-channel whole-head OPM-MEG system is constructed
- System evaluated via repeat measurements of alpha, beta and gamma oscillations
- Two OPM-helmet designs are contrasted, a flexible (EEG-like) cap and a rigid helmet
- The rigid helmet offers significant advantages for a viable OPM-MEG device
- 49-channel OPM-MEG offers performance comparable to established cryogenic devices

## 1. INTRODUCTION

Magnetoencephalography (MEG) (Cohen, 1968) involves measurement of the small magnetic fields generated outside the head by current flow in the brain. Post-measurement modelling of these fields enables construction of 3-dimensional images showing moment-to-moment changes in brain activity. Because MEG offers direct inference on brain electrophysiology, its temporal precision is excellent. Further, in contrast to the potentials measured in electroencephalography (EEG), magnetic fields are relatively unaffected by the inhomogeneous conductivity profile of the head (Baillet, 2017; Boto et al., 2019), meaning that spatial resolution is good (∼5 mm) (Barratt et al., 2018). MEG therefore offers a powerful means to characterise brain function in health and disease. It can be used to assess the formation and dissolution of brain networks in real time as they are modulated in support of cognitive tasks (e.g. Baker et al., 2014; O’Neill et al., 2017) and this has led to its use in cutting edge neuroscience (e.g. Liu et al., 2019). In addition, MEG has potential for clinical application; it is an established tool in epilepsy (e.g. (Stefan and Trinka, 2017)), it has potential for diagnosing disorders like mild traumatic brain injury (Dunkley et al., 2015; Huang et al., 2014) and Autism Spectrum Disorder (Roberts et al., 2019b); and plays an important role in understanding many other disorders, with examples including unipolar depression (Nugent et al., 2015), psychosis (Robson et al., 2016) and dementia (López-Sanz et al., 2018).

Despite this potential, there are a number of drawbacks to the current generation of MEG technology. Detection of the femtotesla-scale magnetic fields generated by the brain is typically made possible via the use of superconducting quantum interference devices (SQUIDs) (Cohen, 1972). However, SQUIDs require cryogenic cooling and for this reason, MEG sensors must be embedded in a cryogenic dewar. This brings about severe limitations: first, the sensor array is fixed in location inside a ‘one-size-fits-all’ helmet. There is (at minimum) a 1.5–2 cm gap between the sensors and the head because a thermally insulating vacuum must be maintained between the participant’s head and the sensors. This gap is inhomogeneous (with the largest brain-to-sensor distances typically in frontal areas (Coquelet et al., 2020)) because the array can’t adapt to different head shapes/sizes. The gap also increases dramatically for individuals with small heads. Since the MEG signal follows an inverse square law with distance, this means poor signal quality for specific brain regions (e.g. frontal lobes), or sometimes across the whole brain (e.g. in babies or children). The second major limitation relates to subject movement; any movement relative to the static sensor array will degrade data quality. Even movements of order 5 mm can be problematic (Gross et al., 2013) and this makes the MEG environment poorly tolerated by many individuals, particularly those with illness inducing involuntary movements such as Tourette’s Syndrome or Parkinson’s Disease. Finally, the cryogenic infrastructure surrounding a MEG system makes both purchase and running costs high, which limits uptake of MEG as an imaging modality.

In recent years there has been significant progress on the development of new magnetic field sensors which have the potential to lift many of the limitations of the current generation of MEG devices. Optically-pumped magnetometers (OPMs) exploit the quantum mechanical properties of alkali atoms to measure small magnetic fields (see Tierney et al., 2019a for a review). OPMs have been shown to have sensitivities close to that of commercial SQUIDs (Allred et al., 2002; Dang et al., 2010; Kominis et al., 2003) and microfabrication techniques have enabled miniaturisation (Griffith et al., 2010; Schwindt et al., 2007; Shah et al., 2007; Shah and Romalis, 2009; Shah and Wakai, 2013) such that OPM packages are now compact. Their potential for measurement of MEG signals has been established (Alem et al., 2014; Johnson et al., 2010; Johnson et al., 2013; Kamada et al., 2015; Sander et al., 2012; Xia et al., 2006), and readily available commercial OPMs now offer a means to develop a new generation of MEG system (Boto et al., 2017). Since OPMs do not require cryogenic cooling they can be placed closer to the scalp than cryogenic sensors, enabling detection of larger signals as well as field patterns with higher spatial frequencies (and consequently higher spatial resolution) (Boto et al., 2016; Boto et al., 2019; Iivanainen et al., 2017; Iivanainen et al., 2019b). Flexibility of placement means an OPM array can, in principle, be adapted to any head size (Hill et al., 2019). In addition, when background fields are controlled (Holmes et al., 2018; Holmes et al., 2019) it is feasible to collect data whilst a participant moves (Boto et al., 2018; Hill et al., 2019). It is therefore possible that the coming years could see a shift in MEG technology, away from fixed cryogenic systems and towards wearable, adaptable, motion-robust systems which provide high quality data. Such a shift would undoubtedly prove a step change for MEG technology, offering access to new subject cohorts (e.g. infants) and new experimental paradigms where head movement is not only allowed, but also encouraged (e.g. Hill et al., 2019; Roberts et al., 2019a).

Although there is exciting potential, OPM-MEG is a nascent technology with significant development still required. Whilst multi-channel systems are available (e.g. (Borna et al., 2017; Boto et al., 2018; Iivanainen et al., 2019b)), most demonstrations still employ small numbers of sensors sited over specific brain regions and the introduction of a whole-head array will be an important step forward. Sensor array coverage (i.e. where to place OPMs to cover all possible cortical locations) and the number of OPMs required to gain parity of performance with cryogenic systems remain open questions (Iivanainen et al., 2019a; Tierney et al., 2019b) and there are, to date, few studies comparing OPM and SQUID measurements. Such comparisons are critical if the MEG community is to gain confidence and adopt OPM technology. Further, whilst the size and weight of OPMs are now appropriate for scalp mounting, the design and fabrication methods for helmets are not established; to date, wearable MEG demonstrations have tended to use 3D-printed helmets, fabricated to fit individual participants (Barry et al., 2019; Boto et al., 2018; Boto et al., 2017; Lin et al., 2019; Tierney et al., 2018). However, such individualised “sensor-casts” are expensive to build and the development of lightweight ergonomic helmets, able to accommodate multiple individuals, would therefore be an important step.

In this paper, we introduce a ‘whole-head’ (49-channel) wearable OPM-MEG system, constructed using commercially-available sensors (QuSpin Inc.). We use this instrument to measure electrophysiological responses to a visuo-motor paradigm. Employing a ‘test-re-test’ experimental design in two participants (each scanned 18 times) we compare the reliability of OPM-measured magnetic field data, and source-space (beamformer-reconstructed) functional images, to an established state-of-the-art SQUID system. We also contrast two different OPM helmet designs, a flexible (EEG style) cap, and an additively-manufactured, generic, rigid sensor-cast. We introduce and evaluate new optical techniques for co-registration of sensor location to brain anatomy for both helmets, and we contrast the trade-offs between flexibility (which ensures OPMs are close to the scalp) and rigidity (which enables accurate knowledge of sensor locations for source reconstruction).

## 2. MATERIALS AND METHODS

*All data were collected by the authors. All code for analysis was custom written in-house using MATLAB.*

### 2.1. OPM-MEG System Description

We built a whole-head multi-channel OPM-MEG system containing 49, 2^nd^ generation, zero-field magnetometers manufactured by QuSpin Inc. (Colorado, USA). Each sensor is a self-contained unit, of dimensions 12.4 × 16.6 × 24.4 mm^3^, containing a Rb-87 gas vapour within a glass cell, a laser for optical pumping, and on-board electromagnetic coils for nulling the background field within the cell. Precisely how this device measures magnetic field has been dealt with in previous papers and will not be repeated here (for a review see Tierney et al., 2019a). The OPMs were mounted on the participant’s head (see below) and connected, via a 60-cm lightweight (3.3 g·m ^-1^) flex cable, to a back-pack. Thicker cables are then taken from the backpack to the control electronics. Analogue output signals were fed from the OPM electronics to a National Instruments digital acquisition system (DAQ). Although these sensors can measure two orthogonal components of the magnetic field, we only measured the component of the magnetic field that was normal to the scalp surface in these experiments.

The system is contained within a magnetically-shielded room (MSR) designed and built specifically for OPM operation (MuRoom, Magnetic Shields Limited, Kent, UK). This MSR, which comprises 2 mu-metal layers and a single copper layer, is equipped with degaussing coils (Altarev et al., 2015), and this, coupled with its novel design, means that the background static magnetic field can be reduced to ∼1.5nT, with field gradients of less than 2 nT·m^-1^. For comparison, a similar MSR with 2 layers of mu-metal and one layer of aluminium in our institution, based on a design typically used to house cryogenic MEG systems has a typical background field of 30 nT with gradients on the order of 10 nT·m^-1^ (this room is not equipped with degaussing coils). Control of background field for OPM measurements is extremely important. The operational dynamic range of the QuSpin zero-field magnetometers (defined as the maximum change in field before gain errors become >5%) is ∼1.5 nT (Boto et al., 2018). Whilst on-board coils enable cancellation of the magnetic fields within the OPM cell, and consequently operation in higher fields (up to 50 nT), these coil currents are set at the beginning of an experiment and usually left unchanged during the recording. This means any movement of the OPM array (e.g. due to head movement) relative to the background field will alter the fields within the OPMs, and render them inoperable. In an MSR with a background field of 30 nT, a head rotation of around 3 degrees is enough to generate the 1.5 nT required to stop an OPM from working. In our novel MSR, an OPM can be rotated through 360 degrees and still maintain operation.

Even though OPMs remain operational in the low background field inside our MSR, head movement within this field still generates artefactual signals which can distort measured brain activity. For this reason, background field and gradients were further controlled using a set of bi-planar coils placed either side of the participant (Holmes et al., 2018; Holmes et al., 2019). These coils, which are wound on two 1.6 m square planes separated by a 1.5 m gap in which the participant is placed, generate 3 orthogonal magnetic fields and linear gradients within a (hypothetical) 40 cm cube inside which the participant’s head is positioned. A reference array, placed behind the participant, then measures the background field/gradient and currents are applied to the bi-planar coils to cancel this remnant field. This takes the background field from 1.5 nT to ∼0.5 nT, which enables a 3 fold improvement in suppression of movement artefacts.

A schematic diagram of the system is shown in Figure 1A. The participant sat on a non-magnetic chair placed in the centre of the MSR between the bi-planar coils (figure 1B). Three separate computers controlled the OPMs, data acquisition, and the stimulus presentation. Note that all control electronics is kept outside the MSR in order to minimise the effect of magnetic interference on the MEG measurements.

**Figure 1:**
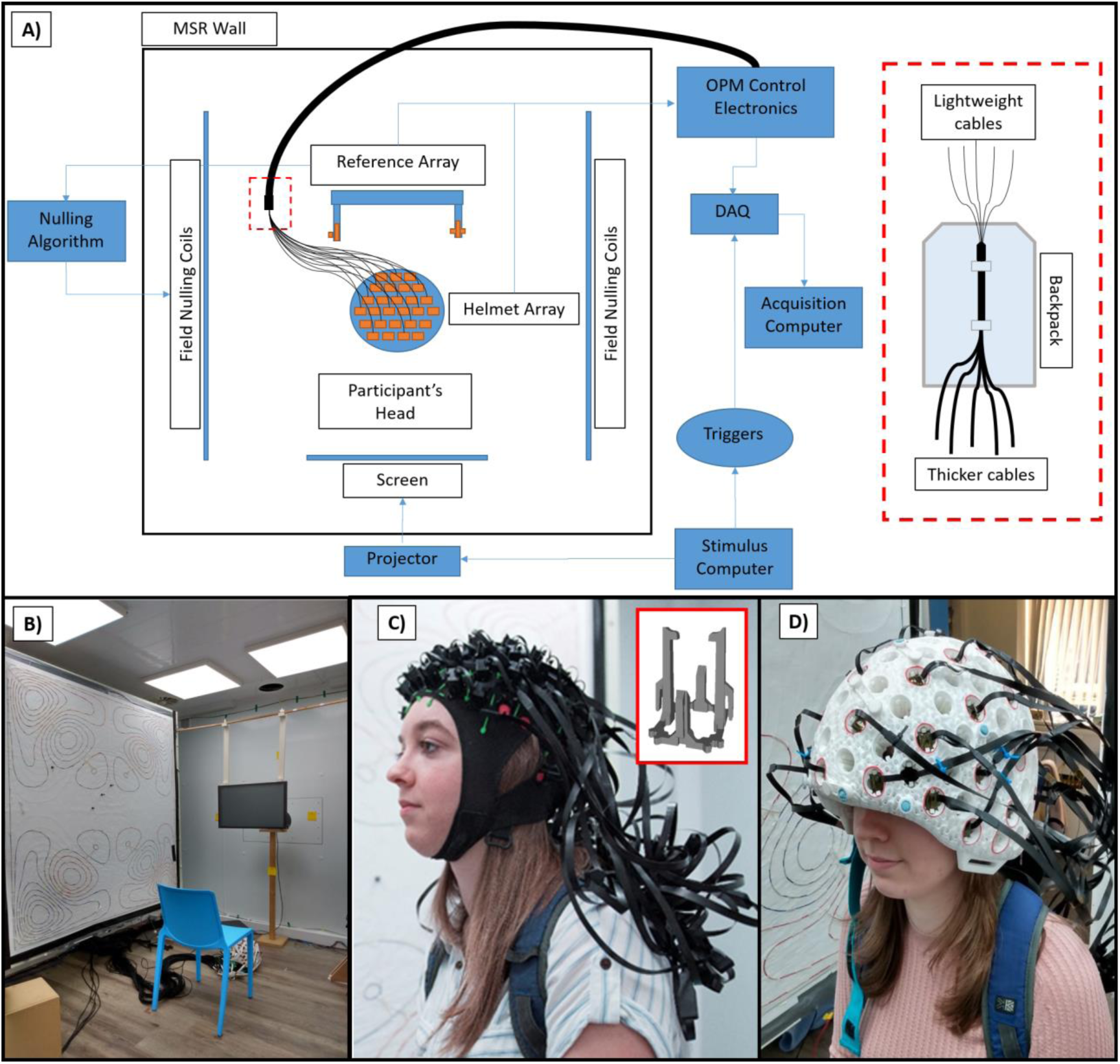
The OPM-MEG system. A) Schematic diagram of the whole system. B) Magnetically shielded room. C) Flexible (EEG style) cap. Both helmets contain push-fit clips to house the 2^nd^ generation QuSpin OPMs (shown inset). D) Rigid additively manufactured helmet.

### 2.2. Helmet Design

Critical to the ultimate design of a viable OPM-based MEG system is the way in which OPM sensors are mounted on the head. This design must represent a balance of four critical considerations: first, sensors must be sited close to the head, to pick up maximum signal, and rigidly held in position (i.e. no movement relative to the head) to avoid artefact. Second, we need accurate knowledge of the sensor locations and orientations relative to each other, and relative to brain anatomy – this is imperative for data modelling. Third, the helmet must be ergonomic and practical (for both participant and operator). Finally, since the current commercially-available sensors require heating of the Rb-87 vapour cell in order to operate in the spin exchange relaxation-free (SERF) regime, the helmet design should allow heat to escape from the OPM and its mounting. Here, we employed two contrasting solutions to this problem:

#### Flexible cap

This is based on an ‘EEG-style’ cap, which contains 63 sensor mounts (see Figure 1C), and is manufactured by QuSpin. The cap is made from elasticated fabric, which is given structure via incorporation of boning. The boning comprises a number of rigid plastic wires which are sewn into the cap to help maintain its shape, and to limit OPM movement relative to the scalp. Whist this gives some structure, the cap remains easily stretched and so readily adapts to any (adult) head shape (it would be easy to manufacture this cap in several sizes to accommodate other age groups). The sensor mounts were 3D printed to house QuSpin 2^nd^ generation sensors, and made from plastic. They hold the sensors at the corners without enclosing them and so heat is able to escape from the external sensor surfaces. The sensor layout is based on the 10:10 EEG system. Flexibility of the cap ensures that sensors are reasonably well positioned close to the scalp surface and so measure high signal (with the caveat that, for individuals with long hair, their hair can push the sensors off the scalp). The cap is also light-weight and comfortable to wear for the participant, with a total weight of 309 g (when containing 49 sensors). However, the flexibility also means that neither the locations nor orientations of the sensors, both relative to one another and relative to the brain anatomy, are known a-priori and this information must be found by using a co-registration procedure.

#### Rigid helmet

In addition, we built an additively manufactured rigid helmet (Figure 1D) from PA12 (which is a nylon polymer) using an EOS P100 Formiga Laser Sintering machine. The size and shape of the inner surface of the helmet is based on a set of adult MRIs; the scalp surfaces were extracted from 9 MRI scans (co-registered together beforehand) and these surfaces were superimposed, to produce a composite head shape. This surface was grown radially by 1.5 mm to form a helmet inner surface that will accommodate the majority of adults’ heads. Ear pockets were incorporated into the design to improve its wearability and comfort. Understandably, the resulting shape is not a one-size-fits-all solution due to the helmet’s rigidity; for any single individual there is naturally an inhomogeneous gap between the scalp surface and the sensors. This gap was later found to range in size over the scalp surface from 1.9 mm to 26.0 mm (median 7.2 mm) for a representative adult male. Whilst any gap is non-ideal, this was significantly smaller than that for a cryogenic MEG system (13.1 mm to 38.8 mm for the same participant (median 26.7 mm)). Again, in the future it would be possible to build this helmet in multiple sizes to better accommodate the normal range of head sizes.

The rigid helmet contains 133 cylindrical mounts, each of which can hold an OPM sensor rigidly at its corners, so eliminating motion of any sensor relative to all other sensors. Motion of the helmet relative to the head is minimised by the use of internal padding and a chin strap. The cylindrical design left space around each of the sensors which was opened up to increase air circulation, enabling the natural convection of heat away from the sensors and also from the participant’s head. The space between the sensor mounts was filled with a matrix-based gyroid triply periodic minimal surface lattice. This provides a lightweight and mechanically-rigid structure, while also facilitating the natural convection of heat away from the head, allowing the participant to feel more comfortable by enabling the flow of air to the scalp. The helmet has a total weight (including sensors) of 1.7 kg (whilst this is quite heavy, future versions of this prototype could be made lighter – see 4. Discussion). A critical feature here is that, although the sensors are not necessarily positioned as close as possible to the scalp surface, the location and orientation of the sensors relative to the helmet (and each other) are known to a high level of accuracy (sub 1-mm and sub 1-degree), because a complete digital representation of the helmet exists; this significantly simplifies the co-registration problem.

### 2.3. Co-registration

A 3-dimensional optical imaging system (structure IO camera (Occipital Inc., San Francisco, CA, USA) coupled to an Apple iPad, operating with Skanect software) and an anatomical MRI scan, were used for co-registration of OPM sensor locations to brain anatomy (Homölle and Oostenveld, 2019; Zetter et al., 2019). The whole-head MRI scan was generated using a 3 T Philips Ingenia system, running an MPRAGE sequence, at an isotropic spatial resolution of 1 mm. The co-registration procedure, for the two helmet types, was as follows:

#### Flexible cap

Immediately following data acquisition, the OPMs were removed from the sensor holders and coloured markers were added in their place (one per sensor). The structure camera was used to acquire a 3-dimensional digitised surface representing the participant’s scalp and face, along with the markers. A colour-thresholding technique was then used to extract the markers, and in this way the OPM locations on the scalp were derived. OPM orientation was taken to be perpendicular to the scalp surface, at the location of the sensor. A surface matching algorithm then fitted the scalp and face surface from the optical scan to the equivalent surface extracted from the anatomical MRI scan, thus resulting in complete co-registration of the sensor locations/orientations to brain anatomy. This procedure is summarised in Figure 2A. Note that it was not possible to add the coloured markers to the OPMs directly, as the surface to be mapped by the structure camera then became too convoluted. It is for this reason that removal of all OPMs prior to digitisation was necessary.

**Figure 2:**
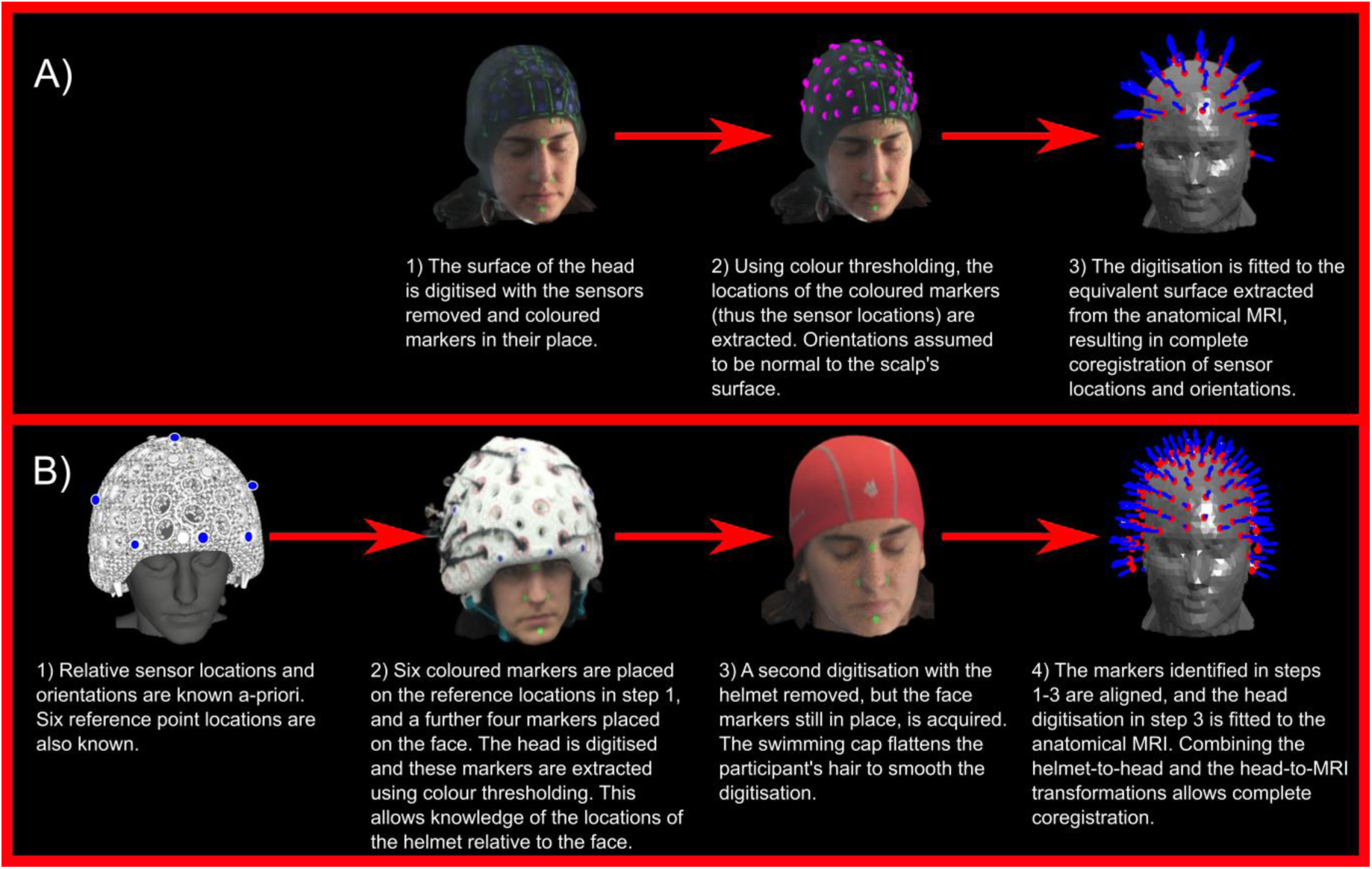
**Schematic diagram showing co-registration algorithm** for A) flexible cap and B) rigid helmet

#### Rigid helmet

For the rigid helmet, the relative locations and orientations of the sensors are known a-priori and this simplifies the procedure, such that it is sufficient to generate a mapping between the helmet and the head (rather than between each sensor and the head). Co-registration was done in two stages. First, 6 coloured markers were placed at known locations on the helmet, and a further 4 were placed on the participant’s face. The camera and colour-thresholding were then used to map the relative locations of these markers, allowing mapping of the helmet to the face. Following this, the helmet was removed and the participant was asked to wear a swimming cap (to flatten down their hair). A second digitisation was acquired of the markers on the face, relative to the rest of the head surface. The head surface was then fitted to the equivalent surface extracted from the anatomical MRI scan. Combining two transforms (helmet-to-head and head-to-MRI) we were able to effect a complete co-registration. This procedure is summarised in Figure 2B.

A key point here is that co-registration error manifests differently in the two cases. For the rigid helmet, any error in co-registration affects all sensors in a similar way – i.e. co-registration error is ‘systematic’-since the relative sensor locations are known very accurately (from the 3D printing process). Conversely, for the flexible cap, we require an independent co-registration of each sensor location and orientation. Co-registration error is therefore different for each sensor and consequently manifests as a ‘random’ error. The consequences of this to source localisation are highlighted by simulations in Appendix 1.

To estimate the accuracy of co-registration, we sequentially placed the two helmets on a single participant, and ran each of our co-registration procedures 10 times. Care was taken to ensure that the helmets did not move between acquisitions. For every sensor, we measured its mean location/orientation (across runs). We then measured the average Euclidean distance from the mean location (across runs) as an estimate of random error in location. Similarly, the average angular difference from the mean orientation was taken as orientation error.

### 2.4. Experimental method

Two participants took part in the study. Both were scanned 18 times; 6 times using OPM-MEG with the flexible helmet (*system 1*), 6 times using OPM-MEG with the rigid helmet (*system 2)*, and 6 times using a cryogenic MEG instrument (*system 3*). Both participants gave written informed consent, and the study was approved by the University of Nottingham Medical School Research Ethics Committee.

#### 2.4.1. Paradigm design

We employed a visuo-motor paradigm which comprised presentation of a visual stimulus which is known to robustly increase the amplitude of gamma oscillations in primary visual cortex (Hoogenboom et al., 2006; Iivanainen et al., 2019c). A single trial comprised 1 s of baseline measurement followed by visual stimulation in the form of a centrally-presented, inwardly-moving, maximum-contrast circular grating (total visual angle 5 degrees; 1.8 cycles per degree). The grating was displayed for a jittered duration of either 1.6 s, 1.7 s or 1.9 s. Each trial ended with a 3 s baseline period, and a total of 100 trials was used. During baseline periods, a fixation dot was shown on the centre of the screen. The participant was instructed to perform abductions of their right index finger for the duration that the stimulus was on the screen, in order to ‘activate’ primary motor cortex (where we expected to see beta-band modulation) as well as visual cortex.

#### 2.4.2. Data acquisition

We acquired data using our OPM-MEG systems with a sampling frequency of 1200 Hz. In the first participant, 42 sensors were available; in the second participant 49 sensors were available. In both cases sensors were spread optimally around the head in order to achieve whole-head coverage (see below). Visual stimuli were presented via back projection through a 21.5 cm diameter cylindrical waveguide in the MSR, onto a screen placed ∼85 cm in front of the participant. We used an Optoma HD39Darbee projector with a refresh rate of 120 Hz. Temporal markers delineating the onset and offset of visual stimulation were also recorded by the digital acquisition system. Co-registration was performed immediately after data collection. Participants were free to move during the scan, but they were not actively encouraged to do so, and movement was not tracked.

Our cryogenic MEG recording used a 275 channel whole-head system (CTF, Coquitlam, BC, Canada), operating in 3^rd^ order gradiometer configuration with a sampling rate of 600 Hz. Note that unlike our OPM system which solely employed magnetometers, the 275 sensors in our cryogenic system comprise 5-cm-baseline, axial gradiometers. These hard-wired gradiometers, coupled with 3^rd^ order synthetic gradiometry, act to reduce background interference in the system, but at a cost of some depth sensitivity in the brain (Vrba and Robinson, 2001). Importantly, the fact that we use magnetometers in our two OPM systems and gradiometers in our cryogenic system makes it difficult to compare sensor-space measurements; this is addressed further below. Again, the stimulus was presented via back projection using a PROPixx (VPixx Technologies) data projector, onto a screen placed 95 cm in front of the participant. For the cryogenic system, co-registration was performed using head localisation coils and digitisation. Specifically, prior to MEG acquisition three head position indicator (HPI) coils were placed at the nasion and pre-auricular points. These coils were energized continuously throughout data collection to allow ongoing assessment of the head location. Following each experiment, a 3D digitiser system (Polhemus, Colchester, Vermont, USA) was used to record a 3D head shape for each participant, which included the locations of the HPI coils. Surface matching of the digitised head shape to a head shape extracted from the participants’ MR images then allowed complete co-registration between brain anatomy and MEG sensor geometry.

#### 2.4.3. Estimating ‘whole head’ coverage

We aimed to estimate the likely coverage of the 3 different MEG systems across the brain. In order to do this, we divided the brain into regular 4-mm voxels. For each voxel, we identified the approximate orientation of the local tangential plane and simulated two dipoles along orthogonal orientations within that plane (i.e. we assumed that radially orientated dipoles would be magnetically ‘silent’). We then calculated the forward field for each dipole using a single-sphere volume conductor model and a dipole approximation (Sarvas, 1987). For both dipoles, we determined the Frobenius norm of the lead-field patterns for each orientation; we then averaged the two values, resulting in an approximation of the total signal strength (across all sensors) from each voxel. This was repeated for all voxels, resulting in a volumetric image showing how estimated signal strength varies across the brain, for any sensor layout. Images were formed based on 49 OPMs in the flexible and rigid helmets and for the conventional cryogenic system. Sensor locations were based on co-registration from a single experiment in Subject 2.

### 2.5. MEG Data analysis

#### Analyses were equivalent for OPM-MEG (with both helmet types) and cryogenic MEG

We first bandpass-filtered all data between 1 and 150 Hz, and removed any ‘bad’ trials (specifically, trials in which the standard deviation of the signal at any one sensor was greater than 3 times the average standard deviation across all trials were removed). In order to visualise sensor space results, data were then further filtered into the beta (13 – 30 Hz) and gamma (55 – 70 Hz) bands (note we also looked in the alpha band (8 – 13 Hz), with results given in Appendix 2). The Hilbert transform of oscillations in each band, for each sensor, was calculated, averaged across trials, and baseline corrected (with baseline calculated over the −3.4 s < t < −2.5 s time window, relative to stimulus offset at t = 0 s). We then measured a sensor-space signal-to-noise ratio (SNR) for all channels: for the gamma band this was calculated as the mean signal in the −2 s < t < 0 s window, divided by the standard deviation of the signal in the 0.5 s < t < 1.5 s window. For the beta band this was calculated as the difference in mean signals in the −2 s < t < 0 s and 0.5 s < t < 1.5 s windows (i.e. the difference between the movement-related beta decrease and the post-movement beta rebound) divided by the standard deviation in the −2 s < t < 0 s window. These SNR metrics were plotted across sensors to visualise the sensor-space topography of the beta and gamma responses. The mean envelope signals for the sensors with the highest SNR were also plotted.

Source localisation was performed using a scalar beamformer (Robinson and Vrba, 1999). The brain was divided into 4-mm cubic voxels and a pseudo-t-statistical approach used to contrast oscillatory power in an active window (−1.2 s < t < 0.3 s) and a control window (0.6 s < t < 2.1 s). Images showing the spatial signature of modulation in oscillatory power were generated for both the beta and gamma bands. Beamformer weights were calculated independently for each band; specifically, covariance matrices were generated using band-limited data and a time window spanning the entire experiment. Covariance matrices were left un-regularised to maximise spatial resolution (Brookes et al., 2008). The source orientation for each voxel was obtained via a search for the highest reconstructed ratio of projected power to noise, and the forward solution was based on a single-sphere model. This algorithm was applied to all 18 experiments in both participants, yielding 18 separate images showing the spatial signature of oscillatory power change, for each band.

Based on the pseudo-t-statistical images, a peak location was determined, and a signal from this location was reconstructed. This was done in two ways: first, using data covariance calculated in the broad (1 – 150 Hz) band, beamformer weights were used to generate a broad-band ‘virtual sensor’ time course. A time-frequency spectrum was then constructed by sequentially frequency filtering the signal into overlapping bands, computing the envelope of oscillatory power, averaging over trials and concatenating in the frequency dimension. Second, using band-limited beamformer weights, the mean envelope signals in the beta and gamma bands were constructed.

Based upon the derived source space images, source space time courses, and sensor-space data, we computed four summary metrics to estimate the effectiveness of each of our three systems:

- **Test-re-test source-localisation consistency:** For each experimental run, we calculated the location of the peak in the pseudo-t-statistical image. These locations were averaged, and standard deviation measured in three orthogonal orientations to define an ellipsoid, spanning the spread of peak locations across runs. The volumes of the ellipsoids were calculated as a summary statistic, with larger values indicating lower consistency of reconstruction.
- **Image consistency:** For each experimental run, the pseudo-t-statistical image was vectorised and Pearson correlation computed between all pairs of runs (i.e. for six runs this results in 15 separate correlation values). These were then averaged and the standard deviation was computed. Here, higher values would indicate greater consistency between runs.
- **Output SNR:** This was calculated based on trial-averaged, beamformer-reconstructed oscillatory envelopes. As for the sensor-space analysis, SNR was defined for the gamma band as the mean signal in the −2 s < t < 0 s window, divided by standard deviation in the 0.5 s < t < 1.5 s window. For the beta band we calculated the difference in mean signal between the −2 s < t < 0 s and 0.5 s < t < 1.5 s windows divided by the standard deviation in the −2 s < t < 0 s window.
- **Output-to-input SNR ratio:** This is effectively a measure of the ability of a beamformer to reconstruct a signal. The measure is simply the SNR of the beamformer-reconstructed time course, divided by the SNR at the best MEG sensor. Values above 1 indicate the beamformer is improving data quality. Importantly, the beamformer not only requires high-fidelity data, but also accurate knowledge of sensor geometry (and consequently accurate co-registration). This simple measure therefore captures the effectiveness of the overall system. However, like input SNR, this is not comparable between cryogenic and OPM systems.

In all cases, summary statistics were computed independently for each frequency band and participant.

## 3. RESULTS

### 3.1. Co-registration accuracy

Figure 3 shows the accuracy of our co-registration procedures for both OPM systems. Recall that sensor locations/orientations were estimated independently 10 times, and averaged. We then calculated the average Euclidean distance of each of the 10 independent sensor locations from the mean, and likewise the average angular orientation difference of each measurement from the mean. This was calculated for each sensor separately [note that these values only provide an estimate of *random* error; i.e. if there was a *systematic* error affecting each run in the same way, it would not be reflected here. However, such an estimate of systematic error would be impossible without a ground truth sensor location/orientation]. In the figure, the upper panels show the results from the rigid helmet and the lower plots show those from the flexible cap; the left-hand plots show error in sensor position whilst the right-hand plots show error in orientation. Quantitatively, results show that for the rigid helmet, the average location and orientation errors (across sensors) were 3.9 mm and 0.94° respectively. The maximum errors at any one sensor were 5 mm and 1.1°, and these tended to occur close to the back of the helmet. This is unsurprising since we used the front of the helmet and the face for co-registration, and small errors at the front of the head will cause rotational inaccuracies which will be amplified at the back of the head. For the flexible cap, the average location and orientation errors were 2.6 mm and 1° and the maximum errors were 3.6 mm / 1°. Here, again, the spatial topographies showed that the largest errors were towards the back of the head. This is likely because the surface matching depends largely on facial features.

**Figure 3:**
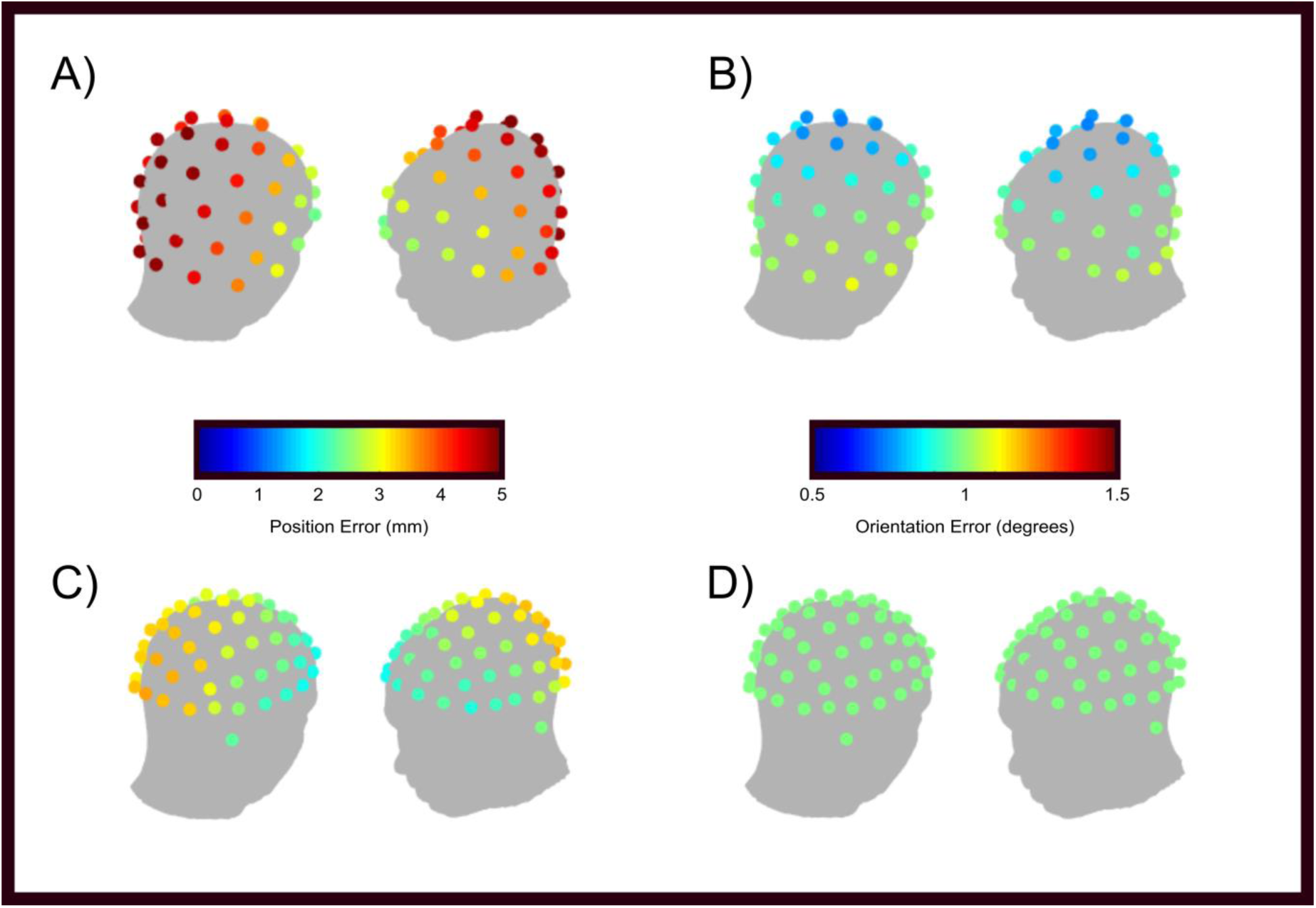
Test-re-test co-registration errors: A) Location error for rigid helmet. B) Orientation error for rigid helmet. C) Location error for flexible helmet. D) Orientation error for flexible cap. Colder colours indicate a more reliable co-registration at that sensor.

Importantly, the magnitude of these errors are relatively small and approximately in line with the likely co-registration errors that are often reported for cryogenic MEG (Adjamian et al., 2004; Chella et al., 2019; Hillebrand and Barnes, 2011; Whalen et al., 2007). We also note that, for the rigid helmet, the error reflects an approximation of how accurately we know the location of the helmet relative to the head/brain since the sensor locations, relative to each other, are known with sub 1 mm and sub 1° accuracy from the additive manufacturing procedure. Conversely, for the flexible cap, whilst the errors are somewhat lower, they reflect how accurately we know sensor positions relative to the brain, and relative to all other sensors. The reader is referred to Appendix 1 for a discussion of the implications of this point.

### 3.2. Sensor array coverage

Figure 4 shows sensor positioning, and coverage in the brain for the rigid helmet (left) the flexible cap (centre) and a cryogenic system (right). In the upper plots, the pink dots show the sensor locations relative to the head surface (though only one aspect is shown, coverage is approximately symmetrical). In the lower plots, the colours represent the norm of the expected magnetic field induced by a unit dipole at all locations in the brain. For all three systems there is a degree of inhomogeneity across the brain with the temporal pole and cerebellum suffering the lowest sensitivity. Nevertheless, we gain reasonable sensitivity over the entire cortex, even with only 49 OPMs. Note the anisotropy of the coverage in the cryogenic system with higher signal strengths in posterior regions and lower signal strengths for the frontal. This is not the case with OPMs in the rigid helmet – indicating a more even coverage. OPMs in the flexible cap also show poor coverage at the front of the head, although this is due to the physical size of the cap and its positioning on the head (to cover visual cortex). Finally note that, as a result of sensor proximity, there is a difference in the scale of the expected fields. Both OPM systems have a marked increase in signal compared to the cryogenic system since no thermally insulating gap is required, and so sensors are positioned closer to the scalp. It’s also noted that the flexible cap gets sensors closer to the scalp than the rigid helmet, hence greater signal is observed.

**Figure 4:**
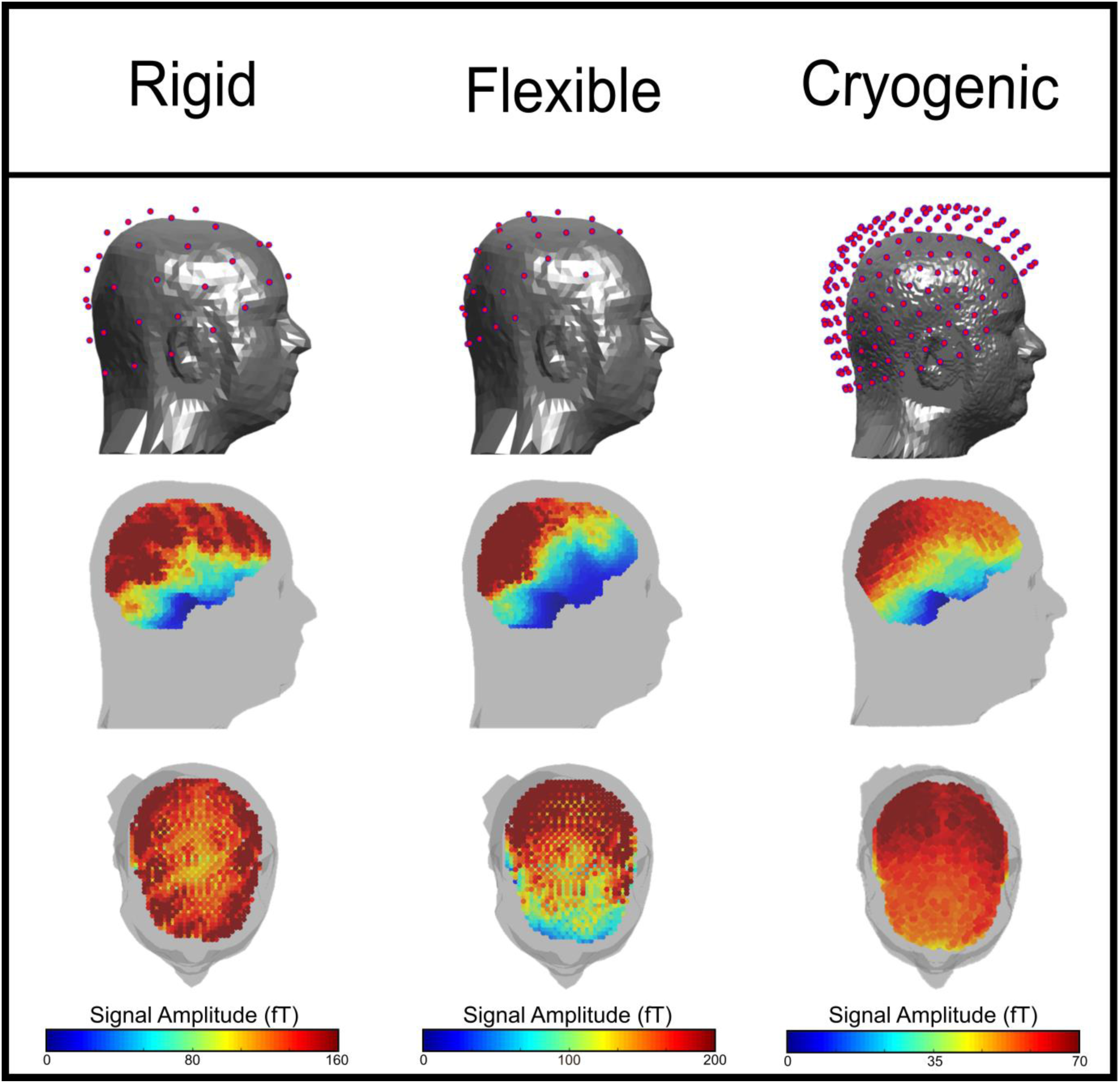
Cortical coverage: The plots on the top row show the sensor locations over the scalp. Lower plots show the norm of the forward fields, for each dipole location in the brain. Left, centre and right shows the rigid helmet, the flexible cap, and the cryogenic system, respectively. Note, the magnitude of the colour axis is different for each system.

### 3.3. MEG data – Sensor-space comparison

Figure 5 shows sensor-space beta- and gamma-band signals. The topography plots show SNR with the six separate cases representing the six repeat measurements. The line plots show the trial-averaged oscillatory envelopes for the beta and gamma bands, extracted from the sensor with the largest SNR (all six runs are overlaid). Importantly, these results show very reliable repeat measurements for all 3 systems – the same sensors show the largest signal change for each run, and the oscillatory envelopes are highly repeatable. Specifically, in the beta band, in sensors over the sensorimotor areas, we observe the characteristic movement-related beta decrease (in the −2 s < t < 0 s window) and the post movement beta rebound on movement cessation (in the 0 s < t < 2 s window). In the gamma band we observe the well-known synchronisation during visual stimulus presentation in sensors over occipital regions. These findings were robust across both participants - see Figure A2 for results in subject 2.

**Figure 5:**
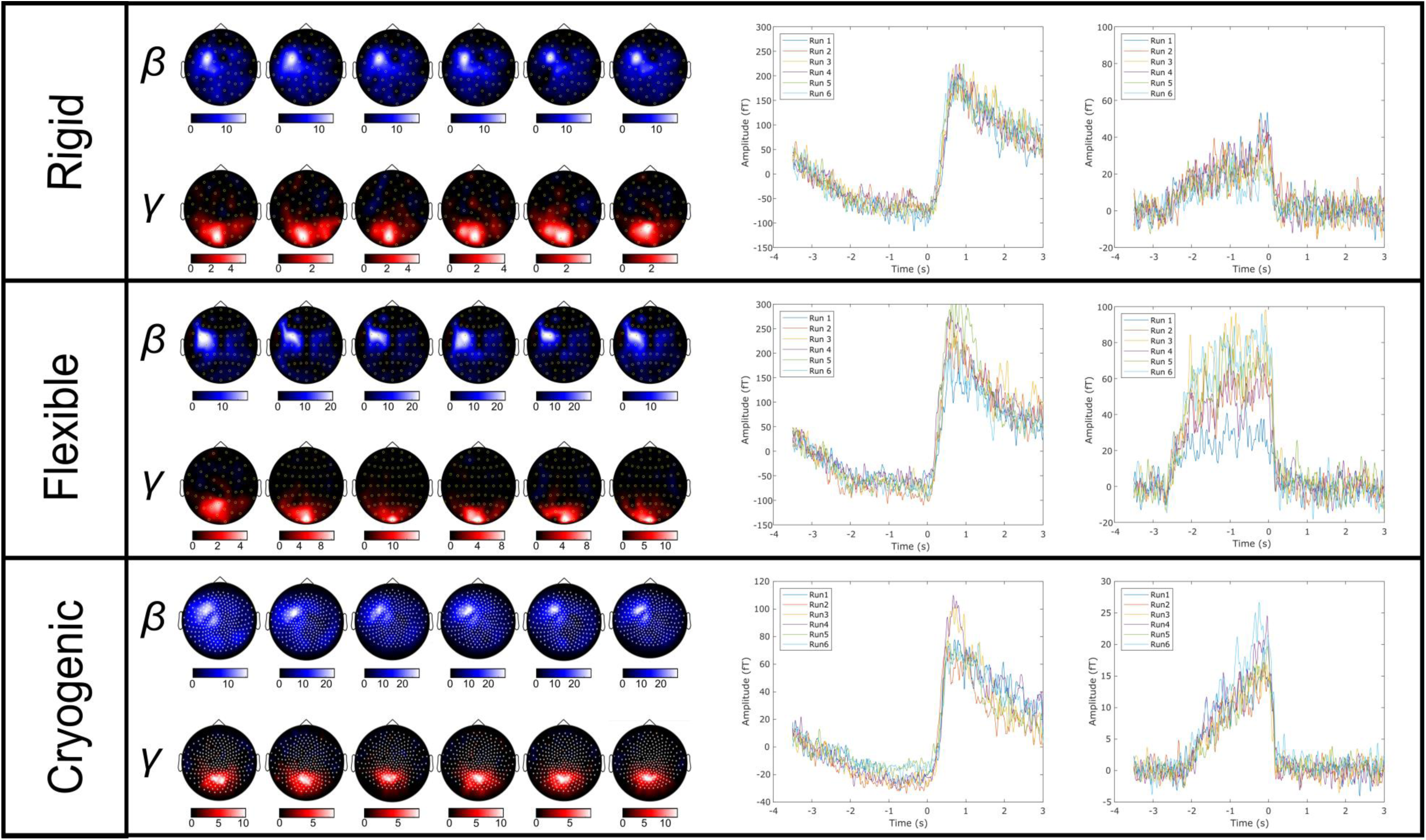
Sensor space results for Subject 1. Upper, middle and lower panels show results from the rigid, flexible, and cryogenic systems respectively. In all cases, the sensor space topography plots show estimated SNRs of the beta and gamma signals for each sensor; the six plots show six repeated measures in Subject 1. The line plots on the right-hand side show the oscillatory envelopes of the beta and gamma effects extracted from the sensor with the largest signal-to-noise ratio (all six runs are overlaid). An equivalent Figure for Subject 2 is shown in Supplementary Material, Figure S1.

Quantitatively, the SNRs are higher for the flexible cap than for the rigid helmet. This is to be expected as the flexible cap holds the sensors closer to the head, on average, than the rigid helmet. The SNR values for the OPMs and cryogenic sensors were comparable, however we note that such values should not, strictly, be compared. This is because cryogenic data are derived from 5-cm baseline axial gradiometers which are processed using third-order synthetic gradiometry based on an independent SQUID reference array; OPM data are completely unprocessed magnetometer data. Given that magnetometers exhibit greater sensitivity to distal sources, it is likely that they show greater contamination from environmental interference sources (including biomagnetic fields from the body and brain regions of no interest). Such environmental sources should be reduced significantly via gradiometry or reference array subtraction. However, the fact that SNR values are comparable despite these differences speaks to the excellent performance of the OPM sensors. This topic will be addressed further in the Discussion.

### 3.4. Source-space comparison and summary statistics

Figure 6 shows the results of source localisation. Figure 6A shows the spatial signature of the change in beta and gamma oscillations for all three systems. Results are for Subject 1, and the equivalent data for Subject 2 are shown in Figure S2. The pseudo-T-statistical images are averaged over all six experimental runs. As expected, for both participants, beta change maps to contralateral primary sensorimotor cortex and gamma modulation originates in primary visual cortex. In Figure 6B, the centre of each ellipsoid represents the mean location of the peak in the relevant pseudo-T-statistical images for beta or gamma modulation. The size of the ellipsoids represents the standard deviation of the peak locations (measured independently in three orthogonal axes). In this way, the ellipsoid volume is a means to estimate the repeatability of source localisation, with lower values indicating a high repeatability. Ellipsoids for the rigid helmet are shown in blue, the flexible cap in yellow, and the cryogenic system in pink. Results for Subjects 1 and 2 are shown in the upper and lower panels, respectively. Note again that the ellipsoids fall close to one another and are well localised to primary sensorimotor and visual regions. Figure 6C displays the ellipsoid volumes for both beta and gamma modulation. The bars in the bar chart show averages (across both subjects) whilst the red and blue squares show the results for each of the two participants separately. Figure 6D shows the image consistency metric (that is, correlation of pseudo-T-statistical images for different pairs of experimental runs, where high values represent high consistency across runs). Again, the bars show average values, whilst the red and blue lines show the case for each participant (the lines are centred on the mean for that participant and the line length shows standard deviation).

**Figure 6:**
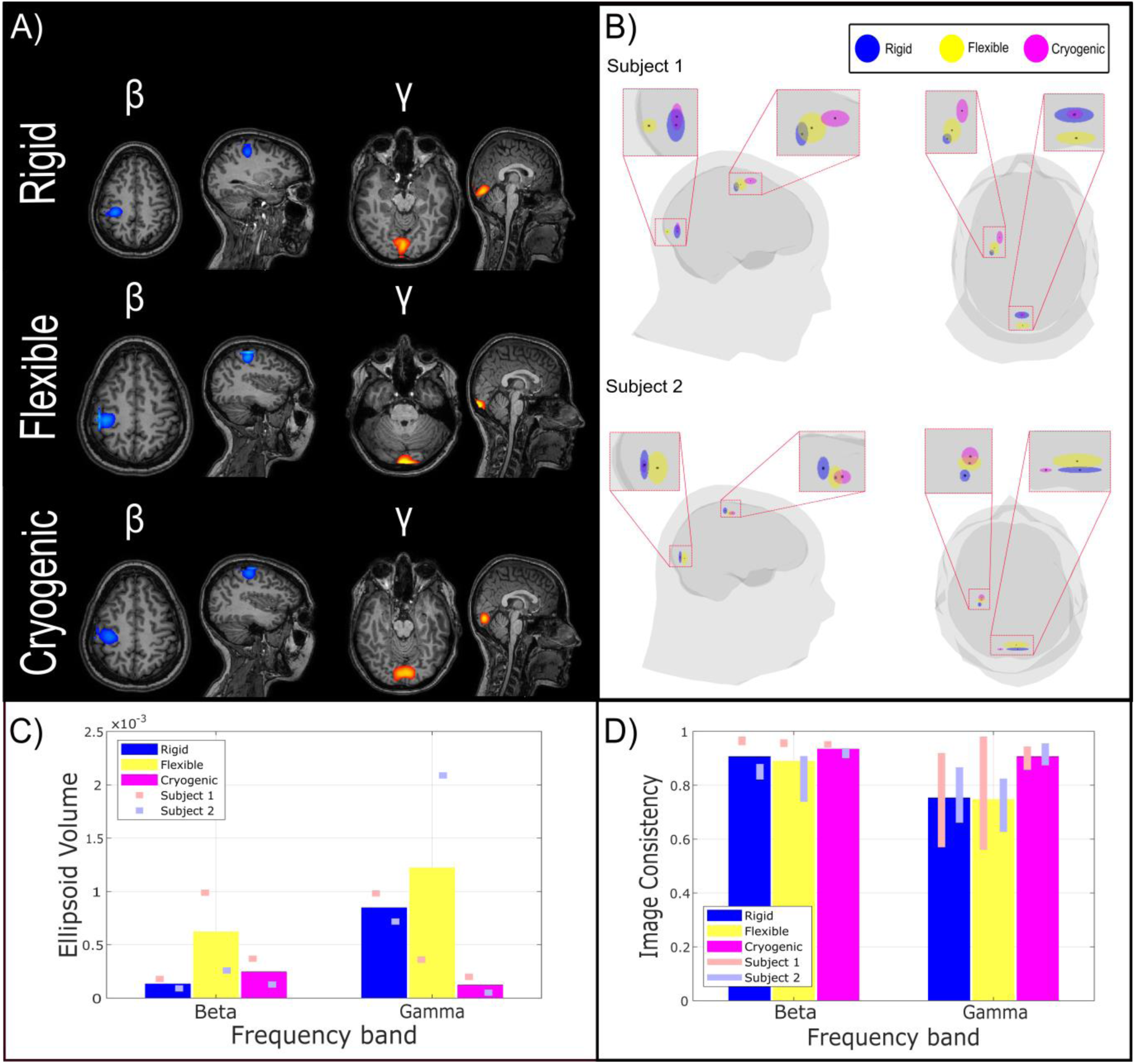
Spatial signature of beta and gamma responses. A) Beamformer pseudo-T-statistical images averaged over all 6 experimental runs for Subject 1. B) Glass brain, with the centre of the ellipsoids showing average peak location across runs. The size of the ellipsoids represents the standard deviation of the peak locations – and hence random variability of localisation across runs. C) Ellipsoid volumes averaged across participants. D) Image consistency (correlation between pseudo-T-statistical images) collapsed across both participants.

Broadly, the repeatability of localisation for all systems is good. Test-re-test localisation results are robustly localised within a small volume, and in all cases correlation between images from separate but identical experiments in the same individual are 75% or better (>90% in beta band). It is noteworthy that these source-localisation metrics characterise both the fidelity of the MEG data, and also the robustness of the co-registration procedure. In the beta band we observe comparable performance for all three systems, whilst for the gamma band, the cryogenic system remains somewhat more reliable. It is also noteworthy that the performance of the rigid helmet appears to be marginally better than the flexible helmet. Appendix A2 shows results for the alpha-band.

Figure 7A shows beamformer-estimated source time courses for the three different systems. The line plots show oscillatory amplitudes in the beta and gamma bands (for all 6 runs overlaid), whilst time-frequency spectrograms (which are averaged over runs) enable a broadband picture of neural oscillatory modulation. The left-hand column shows data extracted from the locations of peak beta modulation (i.e. motor cortex) and the right-hand column shows data extracted from the locations of highest gamma modulation (i.e. visual cortex). Figure 7B shows estimates of the source-reconstructed SNR for the beta and gamma bands. As before, the bars show the average across participants, whilst the lines show the mean and standard deviation for each participant individually. Note that high-fidelity data can be extracted for all three systems, with comparable SNR. The SNR in the gamma frequency band is somewhat higher for the cryogenic system compared with both OPM-based systems, however this is largely a result of lower input (sensor-space) gamma SNR in Subject 2 (see Appendix 2 for alpha results).

**Figure 7:**
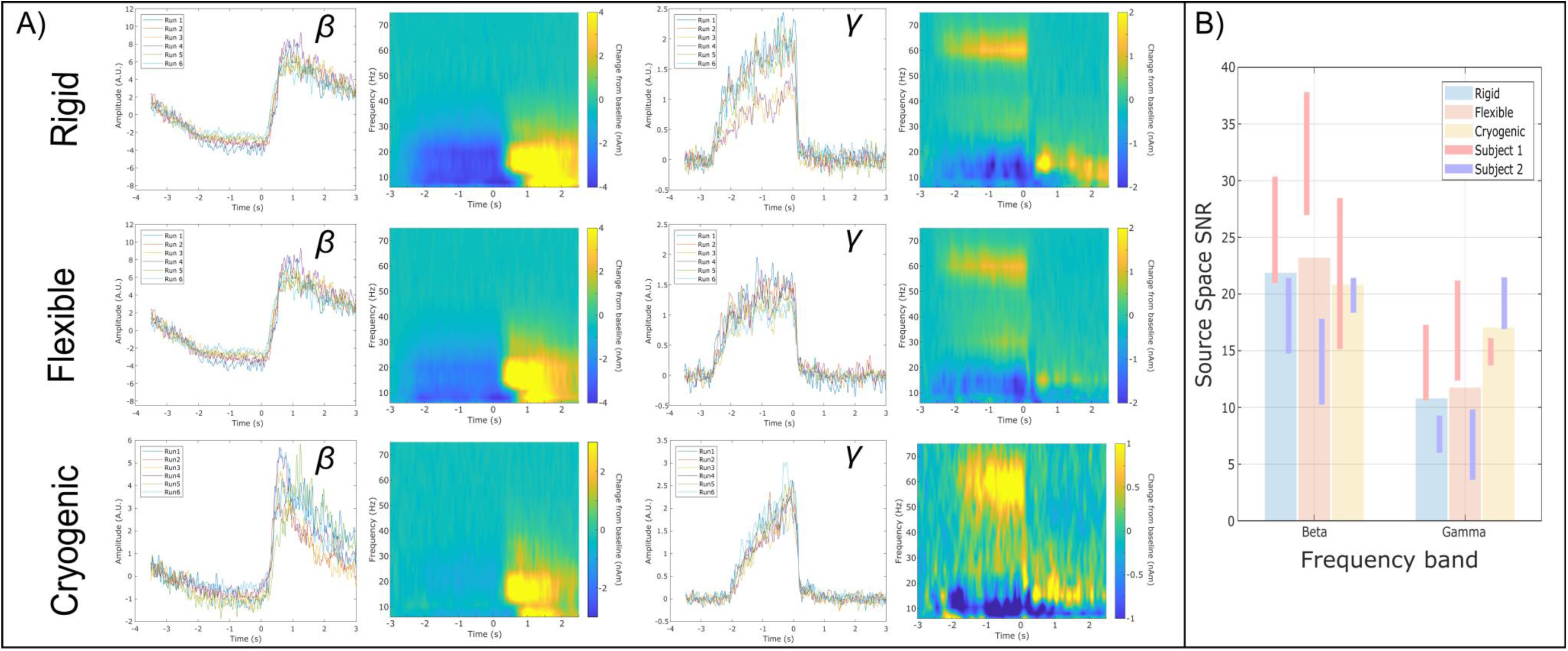
Beamformer-estimated (source space) neural oscillatory activity. A) Oscillatory envelopes and time frequency spectra extracted from the locations of peak beta (left) and gamma (right) modulation. Top, centre and bottom rows show rigid, flexible and cryogenic systems respectively. B) Signal-to-Noise Ratios for the three different systems in the beta and gamma bands.

Finally, Figures 8A and 8B show input (sensor-space) and output (source-space) SNR for the two OPM systems, respectively. In agreement with Figure 3, the sensor-space SNR is larger for the flexible cap than the rigid helmet and this can be attributed to sensors being positioned closer to the scalp surface in the flexible cap and consequently picking up more signal (see also Figure 4). However, in source space, the SNR of the two systems is comparable. Figure 8C quantifies this behaviour by showing the ratio of output to input SNR; values greater than one indicate that application of beamforming is improving data quality by accurate reconstruction, and rejection of signals of no interest that do not originate from the probed brain location. In both systems, beamforming had a positive impact on SNR for both beta- and gamma-band data. However, this impact was more marked in the rigid helmet with e.g. SNR being approximately 2.5 times greater in source space, compared to sensor space, for the gamma band (compared to 1.5 for the flexible cap). This suggests that the beamformer is more effective in the rigid helmet than the flexible cap – such behaviour is in agreement with the results of simulations presented in Appendix 1.

**Figure 8:**
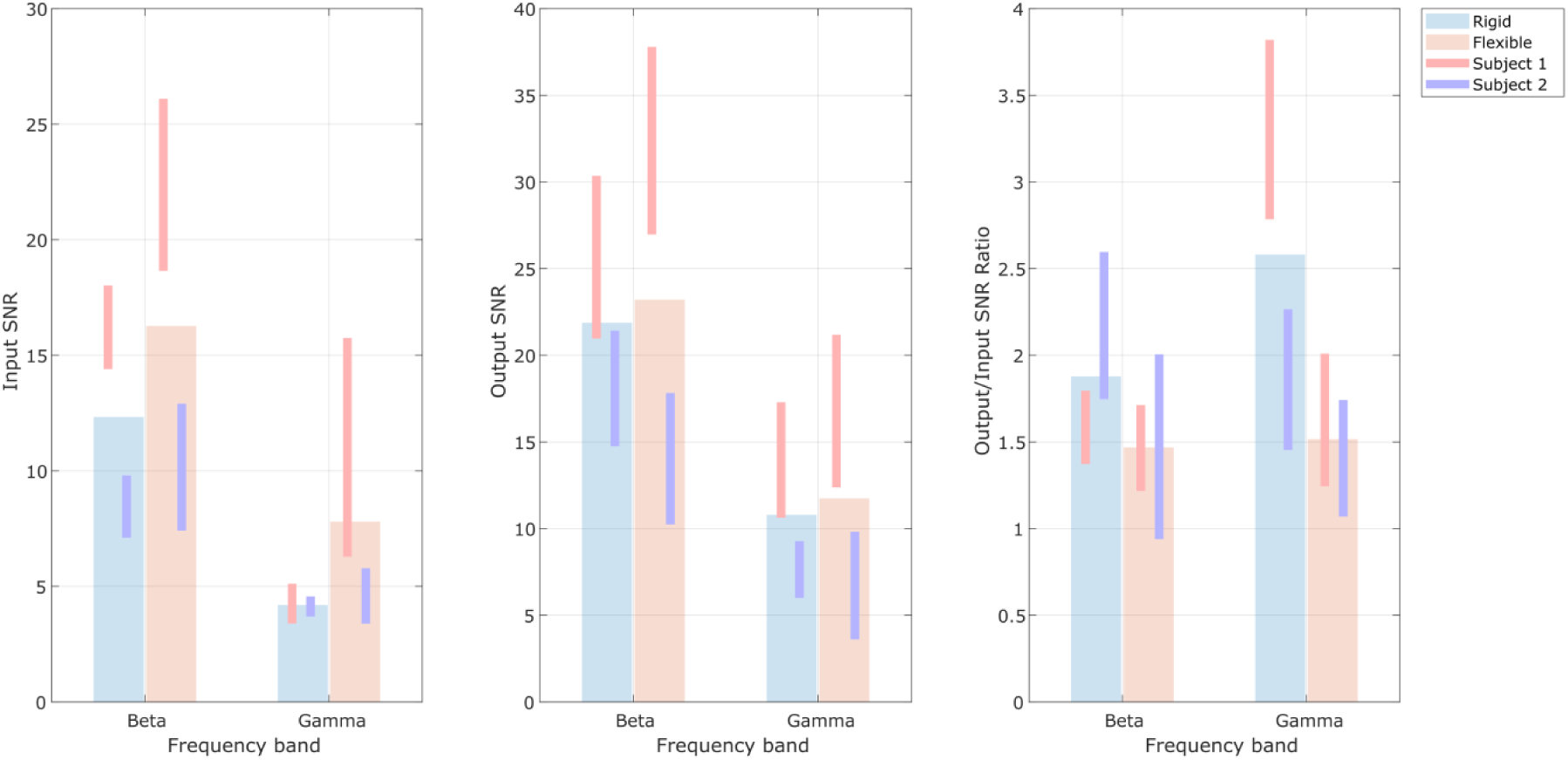
Helmet design comparison: A) Input SNR at the best sensor. B) Output SNR measured in beamformer projected data. C) Ratio of output to input SNR.

## 4. DISCUSSION

We have introduced a 49-channel whole-head wearable MEG system based on commercial OPM sensors. This represents an important step forward compared to previous OPM-MEG demonstrations which have involved smaller sensor counts targeting specific brain regions. The OPMs themselves are second-generation commercial devices – each having a size 12.4 × 16.6 × 24.4 mm^3^, and a weight of 4 g. Previous incarnations of commercial OPMs have been too large, and their cables too heavy, to realistically allow development of a wearable whole–head device (cable weight is a particular problem due to torque on the head). However, these new sensors, with their smaller size and lightweight (3.3 gm^-1^) cabling, represent the ideal building blocks for a complete wearable MEG device. Our current work shows that there are no fundamental barriers to combining large numbers of these sensors (though technical considerations do exist, which are detailed below). The present sensor count (49) remains small compared to cryogenic MEG devices (which have ∼300 sensors). Nevertheless, our array demonstrated better coverage of the cortex than a conventional system (because the scalp-to-sensor distance is more homogeneous around the head). Although some regions (e.g. temporal pole) remain poorly represented, it is likely that, with just a few extra sensors, coverage could be even further improved. Unlike previous demonstrations, in which individualised sensor casts have been used for sensor mounting, here we used generic helmets built to fit many participants – such helmets are ideal for neuroscientific studies where large cohorts are required. We have detailed novel and accurate procedures to coregister these helmets, and consequently sensor location and orientation, to brain anatomy, thus enabling source modelling. Finally, and most importantly, we have demonstrated that in terms of both sensor-space signal detection, and source-space modelling, for alpha- (see Appendix 2), beta- and gamma-band neural oscillations, a ∼50-channel OPM system can offer comparable performance to a current state-of-the-art, cryogenic MEG device.

### 4.1. Helmet design

A primary aim of this paper was to compare two different designs of OPM mounting – a flexible (EEG-style) cap versus a rigid (additively-manufactured) helmet. Both designs are generic, fitting multiple individuals, and both have technical advantages/disadvantages. The sensors are held closer to the head in the flexible cap, and so pick up larger MEG signals than in the rigid helmet, as a consequence of the inverse square relationship between magnetic field and distance from the source. As expected therefore, our results showed that sensor-space SNR (Figure 8A) was consistently higher for the flexible compared to the rigid helmet – particularly for Subject 1 where, for example, gamma band SNR was more than doubled in the flexible helmet. [That said, in Subject 2 gamma SNR was not significantly different between the flexible and rigid helmets, the likely reason is that the shape of the flexible cap was distorted by the participant’s hair, which pushed the sensors away from the scalp over the occipital regions.] In contrast, the greatest (technical) advantage for the rigid helmet is that sensor locations and orientations relative to each other are known accurately. Consequently, any co-registration error affects all sensors in a similar way (i.e. the whole helmet moves) and so this can be thought of as a systematic error. With a flexible cap, co-registration errors are random across sensors. Our simulations (see Appendix 1) show that beamforming is more robust to systematic errors compared to random errors: in both cases one sees an error in the reconstructed location of the source, but with systematic error the fidelity of the reconstructed source time course is significantly improved. This is simply due to a more accurate lead field model – if all sensors are shifted in the same way, the measured field is more likely to look similar to a modelled field, even though the modelled field might appear to originate in the wrong location. If, however, sensor locations are randomly perturbed, the spatial signature of the lead field will be disrupted, and hence will provide a poor match to the measured fields making the beamformer less effective. This finding from simulations is supported by experimental results: the ratio of signal-to-source-space SNR (Figure 8C) is consistently higher for the rigid helmet compared to the flexible helmet, indicating that the beamformer was better able to reconstruct the source accurately. So, whilst the sensor space SNR was greater for the flexible cap, source-space modelling was better when the rigid helmet was used.

There are also practical considerations relating to helmet design. The flexible cap is lightweight (309 g) and adapts to any head size and so provides a comfortable experience for the participant. The rigid helmet is heavier (1.7 kg), more cumbersome, and less adaptable. That said, one advantage of the rigid helmet is that the OPMs are enclosed within a solid structure and therefore protected; future iterations of the helmet could also be made to better manage and protect cables. This all makes the system more robust, and is likely to be extremely useful in subject groups (e.g. infants) who might interfere with the sensors or cables during a scan. It is easy to see how our initial prototype could be made lighter (1.7 kg remains too heavy for effective use in some cohorts), and adaptability could be enabled either via the introduction of multiple helmet sizes, or through the addition of a mechanism to allow sensor travel (in the radial orientation) within the helmet (whilst maintaining accurate knowledge of their location). Another concern is usability. In the flexible cap, we found that all sensors must be removed prior to co-registration. This is because with sensors in place, optical co-registration was ineffective. Removal and replacement of all sensors between experiments was found to take more than an hour. However, for the rigid helmet, sensors can remain in place during co-registration and, consequently, repeat measurements are quicker. Finally, a flexible helmet allows some motion of the OPM sensors relative to each other during a scan (e.g. if a participant moves), whereas a rigid helmet holds them fixed relative to each other. Any movement within a background field generates artefacts. If sensors are rigidly held and helmet motion tracked, those artefacts would be well characterised and, in principle, could be modelled and removed. However, it would be much harder to track independent motion of all sensor locations and orientations. This therefore offers another potential advantage of a rigid helmet.

The findings here indicate that whilst the flexible helmet is comfortable to wear, aesthetically attractive compared to the rigid helmet, and works reasonably well, we believe that the better source-space modelling, ease of use, and system robustness make a rigid helmet a better option for OPM-MEG. The advantage of the rigid helmet would be reinforced by the availability of several helmet sizes or a facility for sensor travel to ensure close proximity of the scalp and the sensor.

### 4.2. Comparison with the existing state of the art

The potential advantages of OPMs over cryogenic MEG systems are well known – adaptability to different head shapes/sizes, motion robustness, no reliance on liquid helium, higher signal strength, higher spatial resolution and lower cost. Not all of these potential advantages have yet been realised, but our data show that an OPM-MEG system, with even a modest sensor count, can compete with the current cryogenic MEG devices. Sensor locations are such that the gap between the scalp and the sensors is more homogeneous across the head (Figure 4). This, in turn, means less variation of sensitivity across the brain. In most adults this likely means improved coverage of the frontal lobes (as in Figure 4), but for people with smaller heads this means better global coverage. Our sensor space data (Figure 5) show the expected increase in signal strength by moving OPMs closer to the scalp compared to cryogenic flux transformers (beta band signals were, on average ∼3 times larger; gamma band signals ∼4 times larger, for Subject 1). However, SNR measures for the SQUID and OPM data are similar, and there are several reasons for this. First, the inherent noise level in our OPMs is higher than in the SQUID sensors used here, which means that whilst moving sensors to the scalp surface affords a signal increase, this is (in part) negated by increased noise. Our previous results from a single (static) OPM show an SNR increase of a factor of two (Boto et al., 2017). However, this assumes sensors touching the scalp and both head and sensor are immobile. Our rigid helmet does not fit perfectly to all participants, and with our flexible cap, participant’s hair distorts the cap shape. This means that the expected increase in SNR compared to a cryogenic device will not necessarily be achieved completely. Second, we have used a wearable device that allows the participant to move freely. Although here the participant was not encouraged to move, slight movement of the head will cause a degree of low amplitude interference. Similarly, at the time of writing there is a known issue with interference caused by relative motion of cables adding to interference (see also below). Both issues can be readily solved by better field nulling (itself a topic of ongoing work) and modifications to sensor electronics. However, the present data were likely negatively impacted by these effects, lowering the OPM-SNR. Finally, as noted above, when comparing sensor-space SNR, we are comparing gradiometer data processed using synthetic gradiometry (cryogenic system), to completely unprocessed magnetometer data (OPMs), with the latter being much more sensitive to interference. With these three considerations taken into account, it is impressive that OPMs show similar performance to the established SQUID-based sensors.

A much better comparison of OPM- and SQUID-based MEG can be achieved via metrics of system performance formed in source space. Here, test-re-test source localisation showed that pseudo-T-statistical images were extremely repeatable for all three systems. Ellipsoid volumes, detailing the spread of peak locations across experimental runs were highly comparable for both the rigid helmet OPM and cryogenic systems, for the beta band. Likewise, our image consistency metric showed better than 90% correlation across runs for all 3 systems. Gamma band results, in both participants, were a little more variable for OPMs compared to the cryogenic system with, on average, larger ellipsoid volumes and lower repeatability. There are likely two reasons for this: first a lower sensor space SNR due to the proximity of the sensors to the head, and second, limited sensor coverage (particularly in the flexible cap) of the visual cortex. Nevertheless, gamma band signals were robustly localised to primary visual cortex and functional images were >75% reproducible.

Perhaps the best measure of MEG system fidelity is source space SNR. This is not plagued by the magnetometer/gradiometer comparison problems that affect sensor space SNR. Moreover, it combines information across channels and requires effective magnetic field modelling, and hence high accuracy of co-registration, as well as high quality input data. In this way it is a good marker of fidelity of the whole system, rather than just of the OPM sensors. Here, again, we showed that source space SNR for the OPM and cryogenic systems were extremely comparable. In the beta band, OPMs showed marginally (but not significantly) improved SNR; in the gamma band, the cryogenic system had better SNR although this was mostly driven by Subject 2; in fact across all experimental runs in Subject 1, gamma band SNR was similar for all three systems. It is important to recognise that source localisation optimally combines information across sensors in a weighted average; it is well known that the more sensors that are available, the better source localisation performs (Boto et al., 2016; Vrba et al., 2004). With this in mind, perhaps the most surprising result of this study is that source localisation is comparable between OPM and cryogenic systems, despite the fact that the cryogenic system has more than 5 times more sensors. However, this appears to support the theoretical findings by Tierney et al., 2019b. It is tempting to speculate that whilst, here, we have shown our OPM system to be “as good” as the current state of the art, as OPM systems inevitably gain more sensors, it is likely that they will significantly overtake cryogenic instruments.

### 4.3. Future challenges

Combining large numbers of OPM sensors into a single MEG system brings about significant technical challenges which should not be underestimated. Perhaps the best examples of this involve some form of ‘cross talk’ between sensors. Firstly, placing two sensors in close proximity affects the measurements made by both sensors. This is a result of magnetic field spread: in order to work, a modulation field – generated by on-board electromagnetic coils within an OPM – is applied to the rubidium vapour. Although this field is strongest inside the cell, it spreads outside the OPM housing and as a result, will penetrate a second OPM in close proximity. This causes changes in both the gain and effective orientation of the sensitive axis of the sensors. Secondly, as noted above, current commercial OPMs operate in the spin exchange relaxation free regime and use a Rb-87 atom vapour, meaning that sensor cells must be heated. In the 2^nd^ Generation QuSpin sensors the cell is electrically heated using an AC current that oscillates at around 400 kHz. Bringing cables in to close proximity to one another, and specifically allowing that proximity to change during a measurement, introduces variable cross-talk between sensors (most likely an effect of capacitive coupling). Thirdly, if heater frequencies are set independently for each sensor, any errors in those frequencies would mean it is possible to pick up ‘beat frequencies’ due to cross-talk between sensors. Finally, as noted above, small movements of sensors within the remnant background field generate low amplitude interference. In the present paper, we addressed some of these concerns – e.g. cross talk between sensors was minimised by ensuring there was a reasonable gap between sensors on the scalp; beating due to cross-talk between heater signals was negated by driving the heaters on all OPMs with synchronised currents of the same frequency. However, most of these concerns mainly affect lower frequencies (e.g. delta band). This is, unfortunately, also where OPMs have an inherently lower performance (noise floor of ∼20 fT/sqrt(Hz) compared to ∼7 fT/sqrt(Hz) for >10 Hz). This means that low frequency measurements are more challenging for an OPM-based system. Nevertheless, improvements to electronics, better cable management, better field nulling, and ultimately improved OPM sensors will undoubtedly meet this challenge.

## 5. CONCLUSION

In conclusion, we have shown that it is possible to construct a ‘whole head’ wearable MEG system based on commercial OPM sensors. We have detailed two different designs for OPM mounting (a flexible cap and rigid helmet) alongside simple and accurate co-registration techniques. Whilst both designs work well, we argue that the rigid helmet is a more judicious choice. Comparing our OPM system to a currently available cryogenic device, our array demonstrated more even coverage, as would be expected. At the sensor level, repeated measurements showed both OPM and cryogenic systems to be extremely reliable. In source space, despite 5 times fewer sensors, our OPM-MEG system showed comparable performance to the established state of the art in terms of source localisation reliability and output (source space) signal to noise ratio, in the alpha (see Appendix 2), beta and gamma bands. OPMs remain a nascent technology and significant technical challenges remain. Nevertheless, this work shows that there are no fundamental barriers to combining large numbers of sensors, and that it is likely OPM-MEG will overtake current systems in terms of performance in the coming years.

## Supporting information

Supplementary Material: Subject 2 Results

## APPENDIX 1: Simulation results

In order investigate how source localisation behaves in the presence of co-registration error, for a rigid helmet and a flexible OPM cap, simulations were undertaken. A single dipolar source was simulated inside a spherical volume conductor (8 cm radius), at a depth of between 2 cm and 2.4 cm from the sphere surface (to approximate a cortical dipole) but otherwise location was randomised. We simulated 49 magnetometers placed 6 mm above the sphere surface and approximately equidistant from each other, covering the upper half of the sphere. The source time course comprised Gaussian random data with standard deviation 1 nAm and this was projected through a forward solution based on a single sphere head model and the dipole equation derived by (Sarvas, 1987). Uncorrelated Gaussian noise was added to each simulated sensor time course with an amplitude of 50fT. A total of 300 s of data were simulated at a sampling frequency of 600 Hz.

We incorporated co-registration errors by both translating the sensor location, and rotating the sensor position and orientation about the origin (which was set as the centre of the sphere). These errors were applied in two ways:

1. **Flexible cap:** To simulate co-registration error using the flexible cap, the error was applied independently to each sensor. The magnitude of the error (e.g. a 1 mm translation and 1 degree rotation) was the same for all sensors, but the direction of these errors were random to simulate the effect of having to determine each sensor location separately via a co-registration procedure.
2. **Rigid helmet:** To simulate the rigid helmet, the same co-registration error was applied to all of the sensors, mimicking the case where the relative locations of the sensors are known accurately, but the error manifests as a shift of the entire helmet on the subjects head.

Note with this model that the magnitude of the errors are the same in both cases; the difference between the two helmet designs is simply in terms of how the error manifests – systematic or random across sensors. We simulated 21 error magnitudes, starting at zero, up to 10 mm and 10 degrees (translation and rotation errors were both applied with equal numerical magnitude in mm and degrees, respectively). Examples are shown in Figure A1A. For each error magnitude the simulation was run for 30 iterations, with new (randomised) dipole location each time

Data were reconstructed using a beamformer. Data covariance was computed in a time frequency window spanning all 300 s and the 1-300Hz frequency range. No regularisation was applied to the resulting matrix. We computed three summary measures in order to estimate the effectiveness of the beamformer:

- **Localisation accuracy:** An image of beamformer-projected source power, normalised by noise (i.e. the pseudo-Z-statistic (Vrba and Robinson, 2001)) was generated within a 12 mm cube, centred on the original simulated source location. The cube was divided into voxels (of isotropic dimension 1 mm) and for each voxel the source orientation was estimated using the direction of maximum signal to noise ratio. We determined the peak location in this image, and computed the displacement between this and the true source location, as a measure of localisation error.
- **Projected power:** This was simply the pseudo-Z-statistic at the peak location in the image. Larger values typically mean a better source estimate (i.e. higher signal to noise ratio).
- **Time course correlation:** At the location of the peak in the beamformer image we calculated the beamformer-projected source time course and measured correlation between this and the simulated (ground truth) time course.

Results are shown in Figure A1B. The left, centre and right panels show the correlation between simulated and reconstructed time courses, pseudo-z-statistic, and localisation error in the beamformer image, respectively. In all cases red shows the rigid helmet and blue shows the flexible helmet. Firstly, we note that co-registration error results in an error in the spatial localisation of the source. This is the case for both rigid and flexible caps, with the flexible cap marginally less prone to this effect. This is likely a result of the randomised (rather than systematic) nature of the error. Increased co-registration error causes a drop in both the projected pseudo-z-statistic and the correlation between simulated and reconstructed time courses. However this drop, whilst apparent for both simulated helmet types, is much more pronounced in the case of the flexible cap compared to the rigid cap (i.e. for the same magnitude of error, we are much more likely to be able to accurately reconstruct the temporal morphology of a source if we use a rigid helmet compared to a flexible cap, albeit with slightly increased localisation error). These findings mirror the experimental results which suggest that the beamformer is more effective in the rigid compared to the flexible cap.

**Figure A1:**
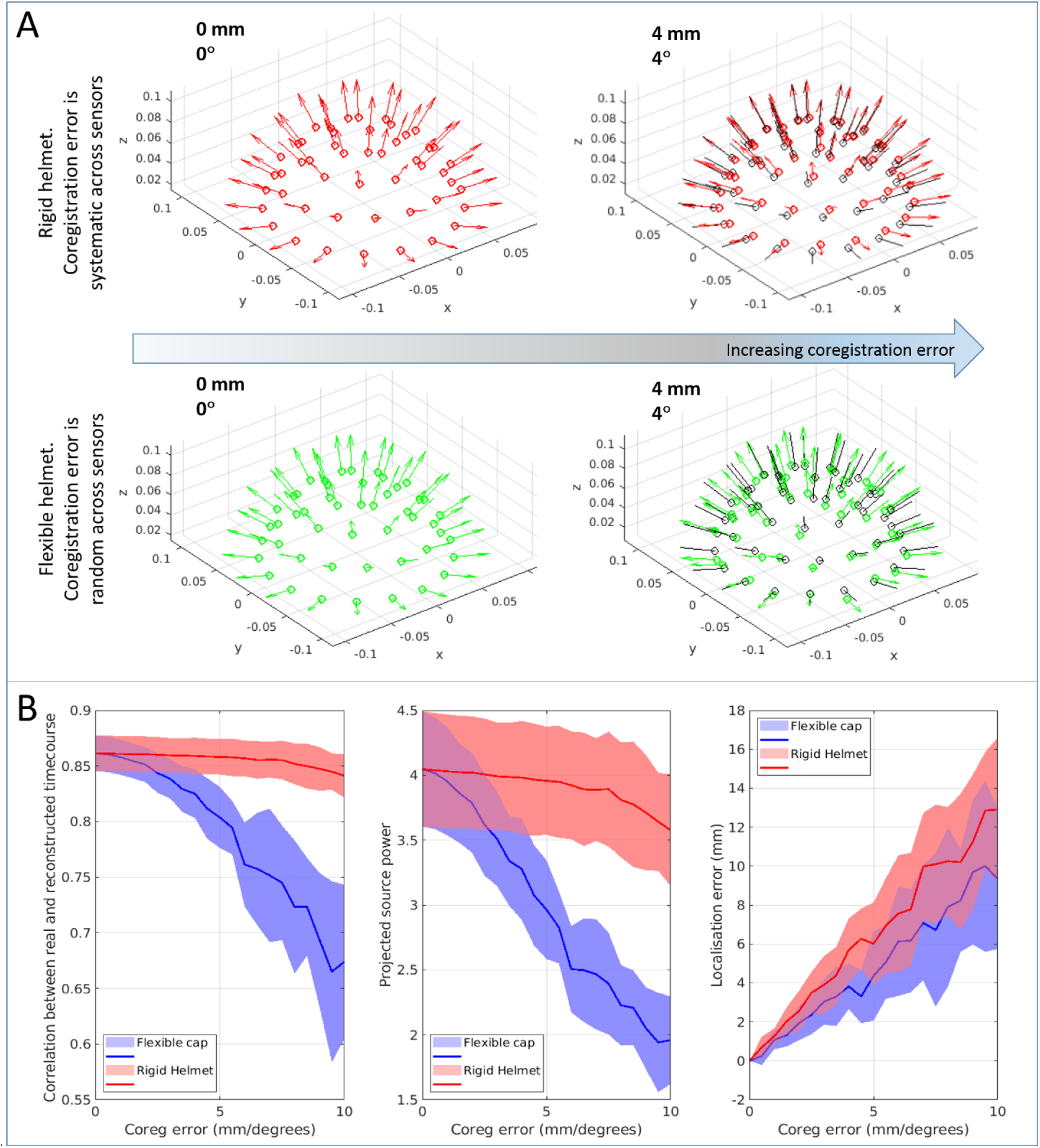
Simulation showing how beamforming is affected by co-registration error in a rigid helmet or flexible cap. *A) Example of co-registration errors in the two cases. For the rigid helmet (upper panel), we assume we know the relative sensor locations and orientations accurately, so co-registration error is systematic across the helmet. For the flexible cap (lower panel), all sensor locations and orientations are acquired from the co-registration process independently, meaning co-registration error is random. The black circles show true sensor positions, and the coloured circles show measured locations with co-registration error. The left hand panel shows zero co-registration error; the right hand panel shows an error of 4 mm translation and a 4 degree rotation (about the origin). B) Summary measures of time course correlation (left) source power (centre) and localisation accuracy (right) plotted as a function of co-registration error. Red shows the rigid cap, blue shows the flexible cap.* Note x-axes represent both translation and rotation – e.g. an error of 5 means 5 mm and 5 degrees.

## APPENDIX 2: Alpha band summary

The same methodology was used to analyse responses in the alpha/mu band (we chose to keep these results in an appendix in order to shorten and simplify the figures main body of our paper). As for the beta and gamma bands, data were first frequency filtered to the 8-13 Hz band, and then processed using a beamformer approach. We derived pseudo-t-statistical images showing the spatial signature of the largest alpha band changes, as well as our summary measures of ellipsoid volume, image consistency, sensor-space SNR and output to input SNR ratio.

Results showed that the largest changes were localised to sensorimotor cortices and probably reflect the well-known ‘mu’ rhythm (an 8-13 Hz) oscillation generated by the sensorimotor system (see Figure A2A). Some alpha changes were also noted over visual cortex. Summary statistics support the findings in the main manuscript: localisations showed that pseudo-t-statistical peaks were largely overlapping for all three systems (Figure A2B), with ellipsoid volumes smallest for the rigid helmet (Figure A2C – left panel). Image consistency (Figure A2C – centre panel) was similar in all three systems for Subject 1, but better in the cryogenic system for Subject 2. Even though sensor space SNR was better for the flexible system (Figure A2D – left panel) source space SNRs were comparable for both the rigid helmet and the flexible cap, and indeed for the cryogenic device (Figure A2C, right panel). As previously, we observed the output/input SNR to be higher for the rigid helmet compared to the flexible cap (Figure A2D – right-hand panel), a result of the improved knowledge of sensor location and orientation. In summary, alpha-band findings strongly support beta and gamma results in showing the advantages of the rigid cap, and moreover that the OPM and cryogenic systems are comparable. This is of some importance due to the known degradation of performance at lower frequency. Specifically, OPM noise levels are higher at low frequency; further, movement artefact and relative cable motion will likely affect the lower frequencies. However, clearly performance of OPM-MEG is not limited down to 8 Hz. The delta and theta bands remain a topic of future research.

**Figure A2:**
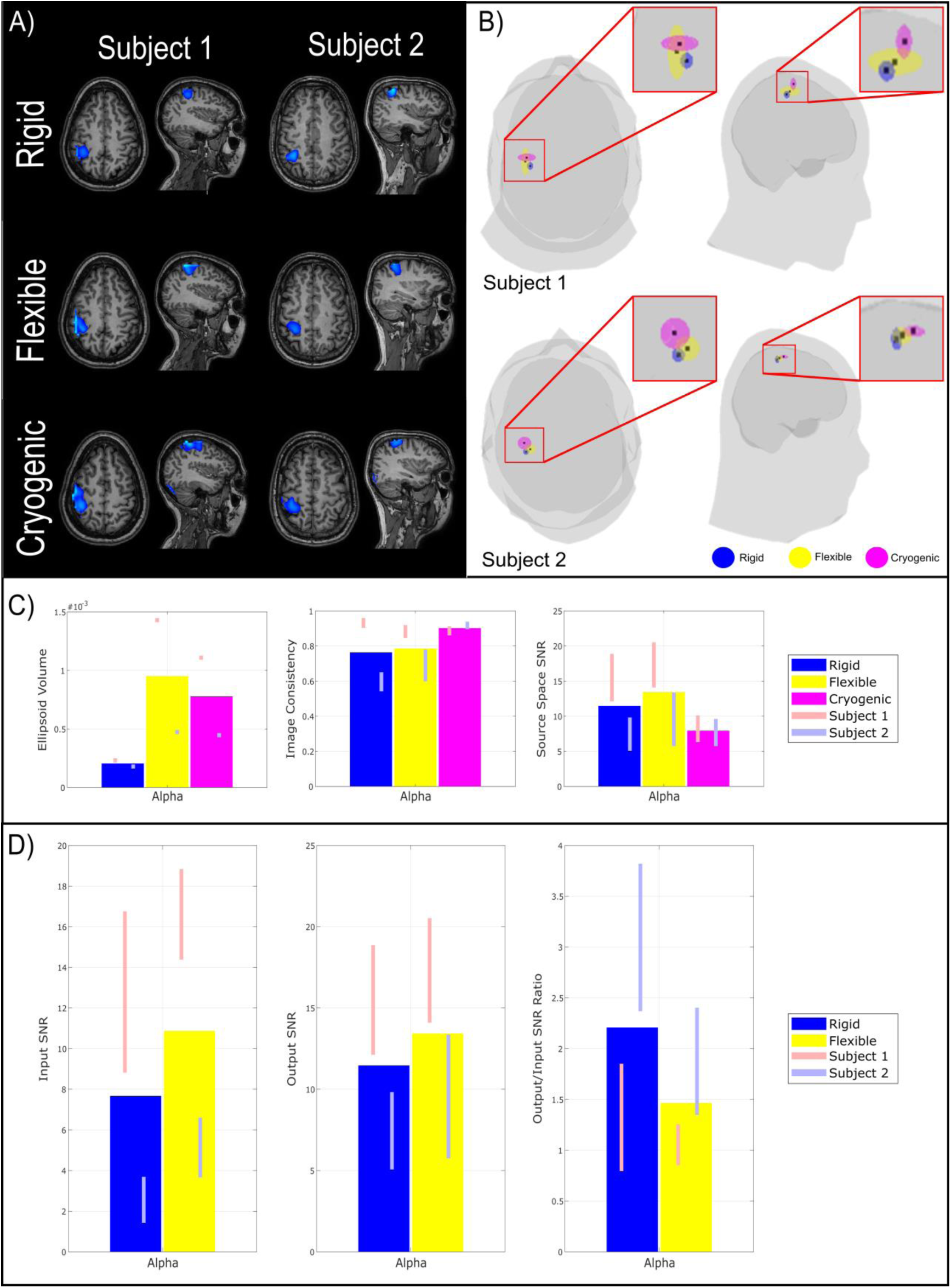
Summary of results in Alpha band. *A) Beamformer pseudo-T-statistical images averaged over all 6 experimental runs for both participants. B) Glass brain, with the centre of the ellipsoids showing average peak location across runs. The size of the ellipsoids represents the standard deviation of the peak locations – and hence variability of localisation across runs. C) (Left) Ellipsoid volumes averaged across participants, (middle) Image consistency (correlation between pseudo-T-statistical images) collapsed across both participants, (right)* Signal-to-Noise ratios for the three different systems in the alpha band. *D) (Left) Input SNR at the best sensor, (middle) Output SNR measured in beamformer projected data, (right) Ratio of output to input SNR.*

## ACKNOWLEDGEMENTS

We express our sincere thanks to collaborators at the Wellcome Centre for Human Neuroimaging, University College London, UK for extremely helpful discussions and ongoing support. This work was supported by the UK Quantum Technology Hub in Sensing and Timing, funded by the Engineering and Physical Sciences Research Council (EPSRC) (EP/T001046/1), and a Wellcome Collaborative Award in Science (203257/Z/16/Z and 203257/B/16/Z) awarded to GRB, RB and MJB.

## CONFLICT OF INTEREST

V.S. is the founding director of QuSpin, the commercial entity selling OPM magnetometers. QuSpin built the sensors used here and advised on the system design and operation, but played no part in the subsequent measurements or data analysis. This work was funded by a Wellcome award which involves a collaboration agreement with QuSpin. All other authors declare no competing interests. Bi-planar coils used for field nulling are available as a product from Magnetic Shields Limited, sold under license from the University of Nottingham.

