## Supplementary Material: Subject 2 Results for "Multi-Channel Whole-Head OPM-MEG: Helmet Design and a Comparison with a Conventional System"

**Supplementary Information: Subject 2 Results**

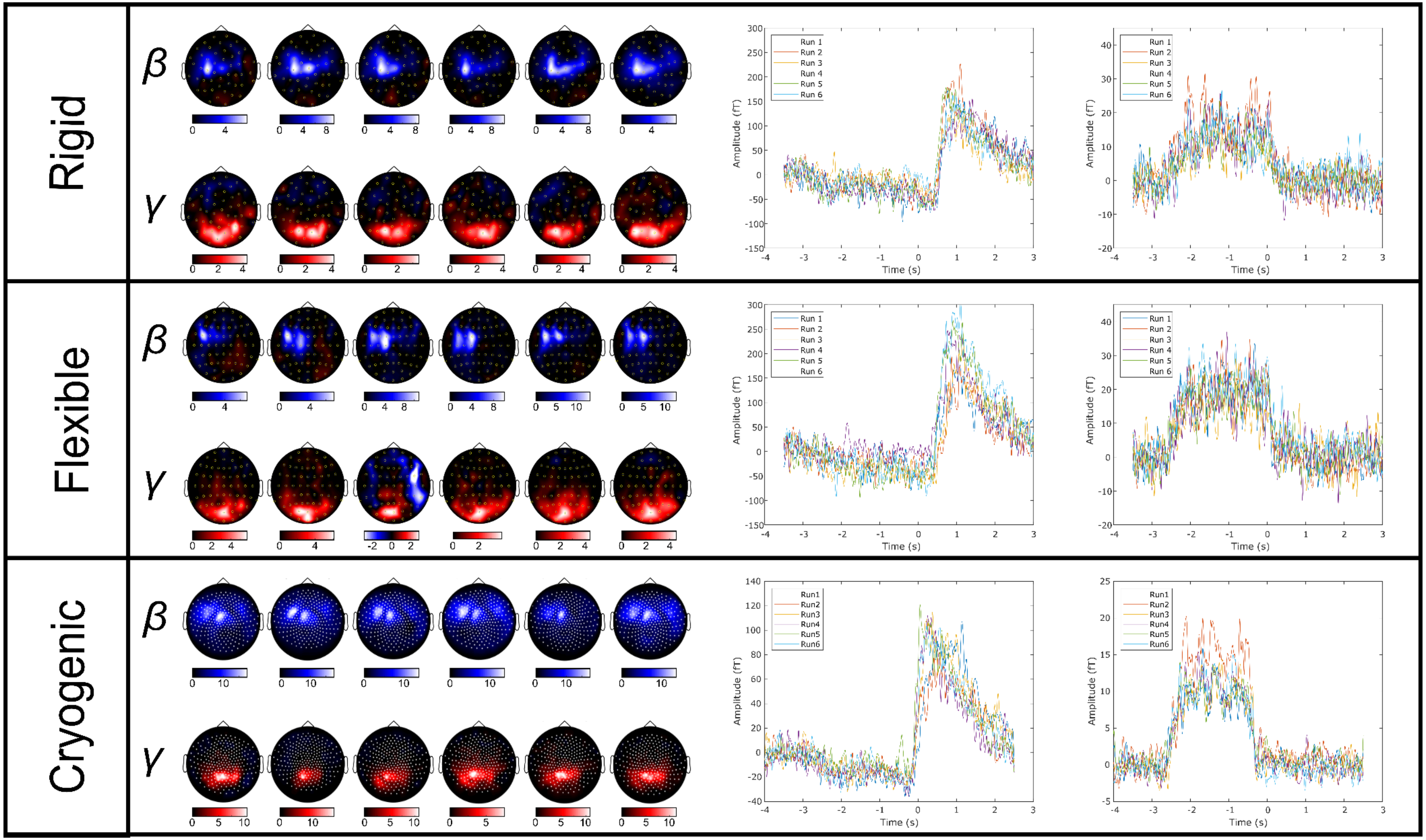


***Figure S1: Sensor space results for Subject 2.*** *Upper, middle and lower panels show rigid, flexible, and cryogenic systems respectively. In all cases, the sensor space topography plots show estimated signal to noise ratios of the beta and gamma signals for each sensor. The line plots show the oscillatory envelopes of the beta and gamma effects.*

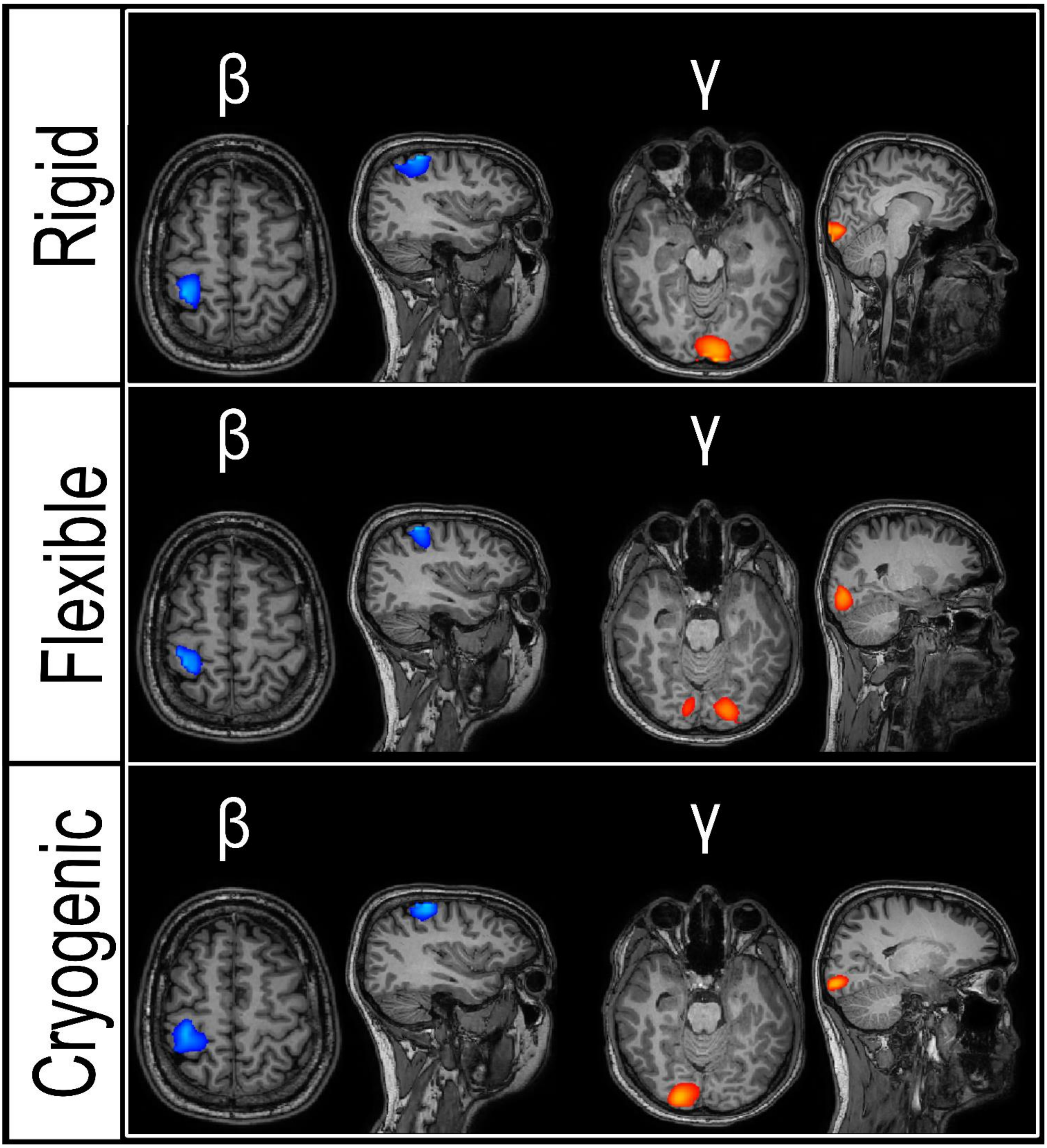


**Figure S2:** ***Beamformer pseudo-T-statistical images averaged over all 6 experimental runs.*** *Results for Subject 2 shown.*

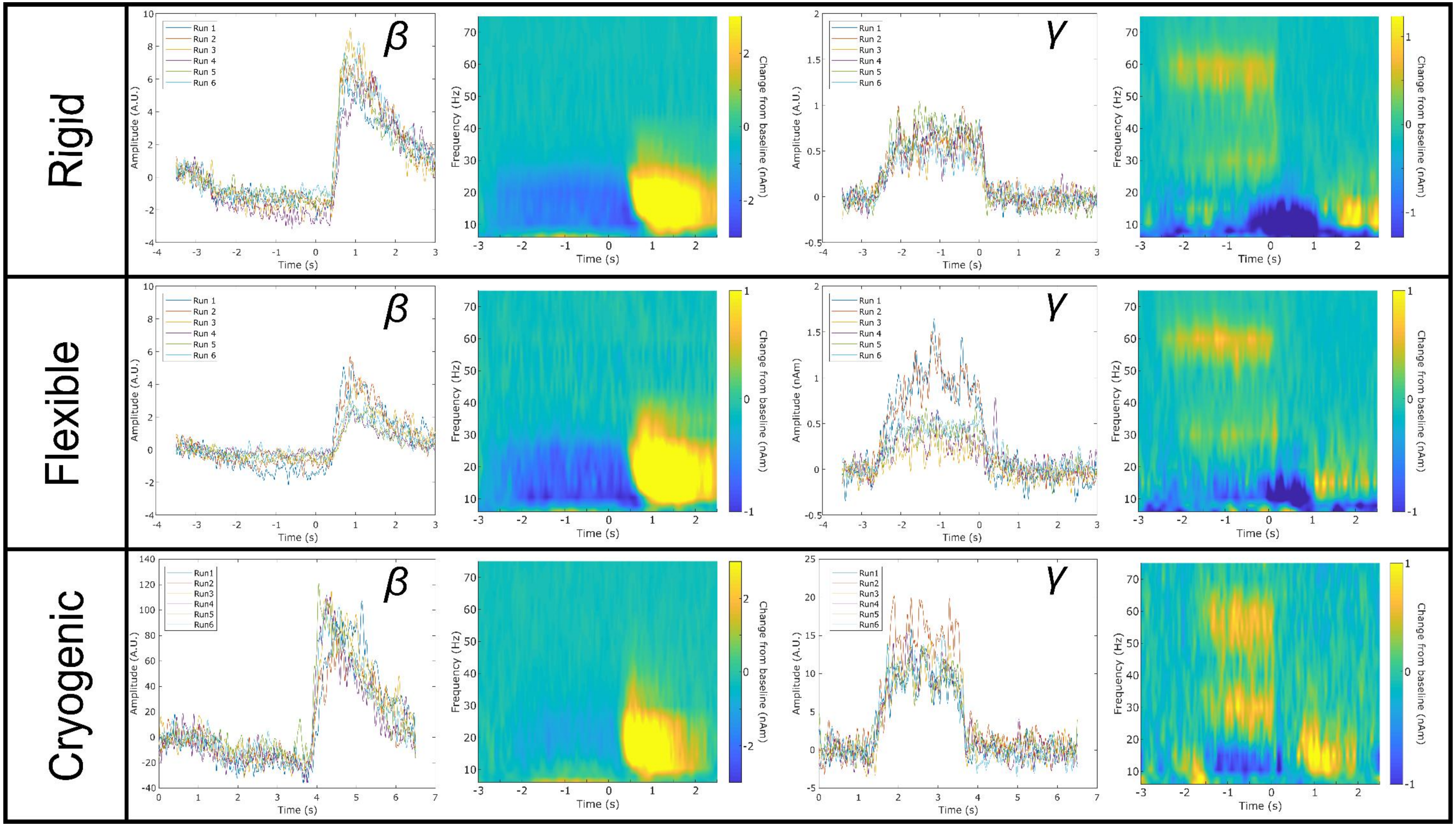


***Figure S3: Beamformer estimated (source space) neural oscillatory activity in Subject 2.*** *Oscillatory envelopes and time frequency spectra extracted from the locations of peak beta (left) and gamma (right) modulation. Top, centre and bottom rows show rigid, flexible and cryogenic systems respectively*
